# Lipid nanoparticle-delivered Cas13b ribonucleoproteins targeting survivin overcome gemcitabine resistance in pancreatic cancer

**DOI:** 10.64898/2026.11.18.767627

**Authors:** Diego Castaño-Esteban, María López-Valls, Mirentxu Santos, Enrique Recuero, Lydia Martín-Vallejo, Demian Pardo, Paola Luciani, Alvaro Somoza, Begoña Sot

## Abstract

Pancreatic ductal adenocarcinoma (PDAC) remains one of the deadliest cancers, largely due to its poor response to chemotherapy and the development of drug resistance. Survivin (BIRC5), an inhibitor of apoptosis frequently overexpressed in PDAC, contributes to tumor cell survival and gemcitabine resistance, making it an attractive therapeutic target. Here, we developed a lipid nanoparticle (LNP)-based platform for the delivery of preassembled pspCas13b ribonucleoproteins (RNPs) to achieve transient and sequence-specific RNA silencing in pancreatic cancer cells. We optimized ionizable lipid formulations for Cas13b RNP encapsulation and identified a formulation achieving approximately 80% encapsulation efficiency, with sustained colloidal stability and gene-silencing activity during seven days of storage at 4 °C. LNP-delivered Cas13b RNPs efficiently silenced endogenous BIRC5 in three pancreatic cancer cell lines, reducing survivin protein levels and inducing apoptosis. In gemcitabine-resistant PANC-1 cells, BIRC5 silencing synergistically enhanced the cytotoxic effect of gemcitabine in vitro. This therapeutic combination was further evaluated in a PANC-1 xenograft model, where only the combined administration of BIRC5-targeting Cas13b RNP-LNPs and gemcitabine significantly inhibited tumor growth, without apparent systemic toxicity. Intratumoral administration also restricted nanoparticle distribution to the injected tumors. These findings establish LNP-mediated delivery of Cas13b RNPs as a new strategy for transient RNA-targeted therapy and provide proof of concept for combining survivin silencing by Cas13b with chemotherapy to address gemcitabine resistance in pancreatic cancer.

## 1. Introduction

Pancreatic ductal adenocarcinoma (PDAC) is the third leading cause of cancer death in the United States, with a five-year relative survival of only 13%^1^. Most patients present locally advanced or metastatic disease and receive gemcitabine-based or FOLFIRINOX regimens^2,3^ that provide only modest survival gains. Intrinsic and acquired chemoresistance, reinforced by a dense and poorly perfused stroma that limits drug delivery, is a central cause of treatment failure^4^. New targets and modalities able to overcome resistance are therefore urgently needed.

Survivin (BIRC5) is a member of the inhibitor of apoptosis protein family^5^. It is low or undetectable in most differentiated adult tissues but re-expressed in many tumors, which offers a therapeutic window^6^. In PDAC, survivin is detected in 88–94% of tumors^7^, and its cytoplasmic localization has been associated with shorter survival^8^. Survivin also contributes to gemcitabine resistance. The resistance of PANC-1-derived stem-like cells depended on survivin expression^9^, and gemcitabine itself up-regulates survivin in MIA PaCa-2 cells, whereas survivin silencing by RNA interference increased their sensitivity to the drug^10^. As a small non-enzymatic protein, survivin is difficult to inhibit pharmacologically, and RNA-targeting strategies have therefore been explored. Antisense oligonucleotides reduced survivin levels in some patients, but they did not improve outcomes in randomised phase II trials and caused dose-limiting haematological or hepatic toxicities^11^. These failures have been attributed to insufficient and inconsistent target engagement, poor delivery to tumours and chemistry-related toxicities^5^. A silencing modality with different properties, delivered locally, is thus worth exploring.

CRISPR–Cas13 enzymes are RNA-guided RNases that silence transcripts with high specificity, without altering the genome^12,13^. Cas13 has been used to silence mutant KRAS in pancreatic cancer cells and xenografts^14,15^. The in vivo strategies for Cas13 have, however, relied on sustained expression of the effector, which carries the risk of collateral degradation of bystander RNAs and of toxicity^16,17^. Delivering the preassembled Cas13 ribonucleoprotein (RNP) would make the activity transient, and RNP delivery has been associated with fewer off-target events than plasmid expression for Cas9^18^. Lipid nanoparticles (LNPs) can deliver Cas9 RNPs in vivo^19,20^, whereas Cas13 RNPs have been delivered only with virus-like particles^21^. To our knowledge, they have never been encapsulated in ionizable-lipid LNPs.

Here we developed LNPs for pspCas13b RNPs and used them to silence BIRC5 in pancreatic cancer cells. The nanoparticles induced apoptosis without detectable collateral RNA degradation and acted synergistically with gemcitabine in gemcitabine-resistant PANC-1 cells. When injected intratumorally, only the combination with gemcitabine inhibited the growth of PANC-1 xenografts.

## 2. Results

### 2.1. pspCas13b RNP mediates gene knockdown eukaryotic cells

With Cas9, the use of RNPs has been shown to be safer and more efficient than the use of DNA. However, most studies involving Cas13 have been performed using DNA or mRNA. Therefore, we evaluated the activity of PspCas13b RNPs after direct delivery into eukaryotic cells by lipofection. PspCas13b was selected because of its previously reported high activity in eukaryotic cells and its lower toxicity, attributed to limited collateral activity^22^. GFP knockdown was assessed in HeLa cells constitutively expressing GFP using two guide RNAs, guide 1 and guide 2 (Figure S1).

Only guide 1 produced efficient GFP knockdown, and was thus selected to conduct the following experiments. GFP expression by flow cytometry, qPCR, and western blot was also assessed Fusing guide 1 (Figure 1). GFP reduction of expression was clear using the three techniques.

**Figure 1.**
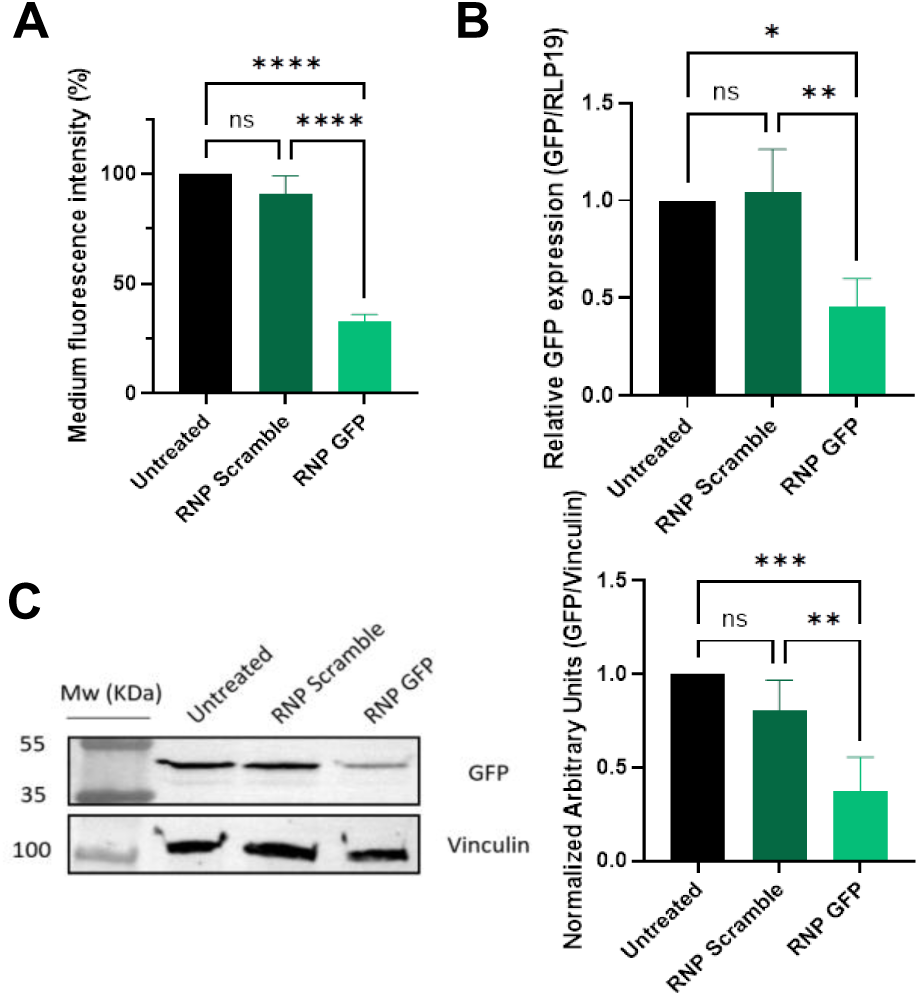
GFP knockdown in HeLa Cells by Cas13b. Cas13b RNP assembled with guide 1 was transfected into HeLa cells by lipofection. GFP knockdown was assessed by flow cytometry (A), qPCR (B) and western blot (C). Western blot signal was quantified using ImageJ and normalized through vinculin expression. Data are presented as mean ± SD from three independent biological replicates (n = 3). Statistical differences were determined by one-way ANOVA in GraphPad Prism. Brackets indicate the groups compared. ns, not significant (P > 0.05); *, P ≤ 0.05; **, P ≤ 0.01; ***, P ≤ 0.001; ****, P ≤ 0.0001.

### 2.2. Optimization of Lipid nanoparticles formulations for the delivery of pspCas13b RNP

Nine different formulations were tested (Table 1), using two different ionizable lipids, several helper lipids, and either including or omitting the cationic lipid DOTAP to enhance Cas13b RNP binding at physiological pH^19,20^. LNPs were prepared by manual mixing: an ethanolic lipid solution was added to an aqueous solution of preassembled PspCas13b RNP (Cas13b:crRNA molar ratio of 1:2) at a 1:3 (v/v) organic:aqueous phase ratio, gently mixed by pipetting, and incubated for 15 min at room temperature (see Materials and Methods).

**Table 1.** Lipid formulations composition.

| **Formulations** | **Composition (molar %)** |
|---|---|
| F0 | C12-200: DOPE: Chol: DMG-PEG: DOTAP (31.5: 14.4: 41.9: 1.4: 10) |
| F1 | DSPC: DOPE: Chol: DMG-PEG: DOTAP (31.5: 14.4: 41.9: 1.4: 10) |
| F2 | C12-200: DSPC: Chol: DMG-PEG: DOTAP (31.5: 14.4: 41.9: 1.4: 10) |
| F3 | C12-200: DOPE: Chol: DMG-PEG: DSPC (31.5: 14.4: 41.9: 1.4: 10) |
| F4 | C12-200: DOPE: Chol: DMG-PEG: DOTAP (50: 5: 38.5: 1.5: 5) |
| F5 | C12-200: DSPC: Chol: DMG-PEG: DOTAP (50: 5: 38.5: 1.5: 5) |
| F6 | DLin-MC3-DMA: DOPE: Chol: DMG-PEG: DOTAP: DSPC (50: 2: 38.5: 1.5: 2:6) |
| F7 | DLin-MC3-DMA: DOPE: Chol: DMG-PEG: DOTAP: DSPC (50: 3: 38.5: 1.5: 3: 4) |
| F8 | DLin-MC3-DMA: DOPE: Chol: DMG-PEG- DOTAP (50: 5: 38.5: 1.5: 5) |
| F9 | DLin-MC3-DMA: DOPE: Chol: DMG-PEG: DOTAP (48.9: 4.9: 37.7: 3.7: 4.9) |

First, the hydrodynamic diameter and polydispersity index (PDI) of the LNP formulations were measured by DLS (Table 2). Formulations with a diameter greater than 250 nm were excluded. The toxicity of the remaining formulations (F0, F3, F4, F5, F6, F8 and F9) was then assessed (Figure S2A). Only F3, F6, F8 and F9 were non-toxic to HeLa cells. Therefore, these three formulations were evaluated for intracellular delivery of Cas13b RNPs and subsequent GFP knockdown (Figure S2B). Only F8 and F9 reduced efficiently GFP fluorescence. F3 lacks the positively charged lipid DOTAP, which may impair its ability to efficiently encapsulate Cas13b RNPs. F6, F8 and F9 have similar compositions, but F8 and F9 contain higher DOTAP concentrations and is therefore expected to interact more effectively with RNPs. F9 was selected for subsequent experiments because it achieved GFP knockdown comparable to that of F8 (Figure S2B) while exhibiting a smaller hydrodynamic diameter (Table 2). These results were confirmed by flow cytometry, qPCR, and western blotting (Figures 2A–C). Furthermore, Cas13b RNP encapsulation was assessed by measuring crRNA encapsulation efficiency (Figure 2D).These findings identify F9 as the most suitable formulation for Cas13b RNP delivery and GFP knockdown in HeLa cells.

**Figure 2.**
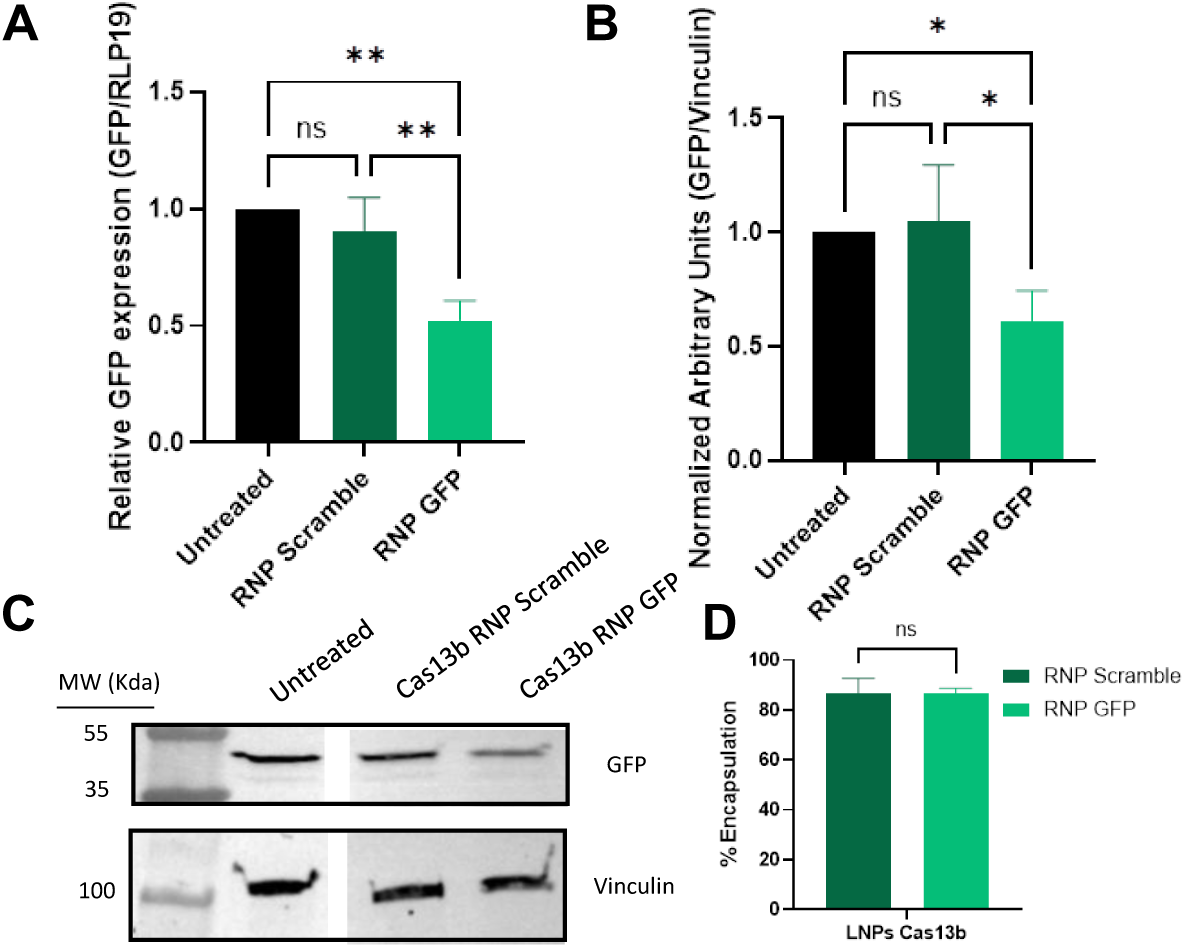
Knock down activity of Cas13b encapsulated into LNPs. A) qPCR of GFP mRNA in Hela cells treated with LNPs. (n=3). B) WB experiment to assess the decrease of GFP expression after treatment (n=3) C) WB signal quantification (n=3). An scramble crRNA was used as control. D) Encapsulation efficiency of Cas13 crRNA into LNPs measured by qubit fluorescence. Data are presented as mean ± SD from three independent biological replicates (n = 3). Statistical differences were determined by one-way ANOVA or t-test (D) in GraphPad Prism. Brackets indicate the groups compared. ns, not significant (P > 0.05); *, P ≤ 0.05; **, P ≤ 0.01; ***, P ≤ 0.001; ****, P ≤ 0.0001.

**Table 2.**
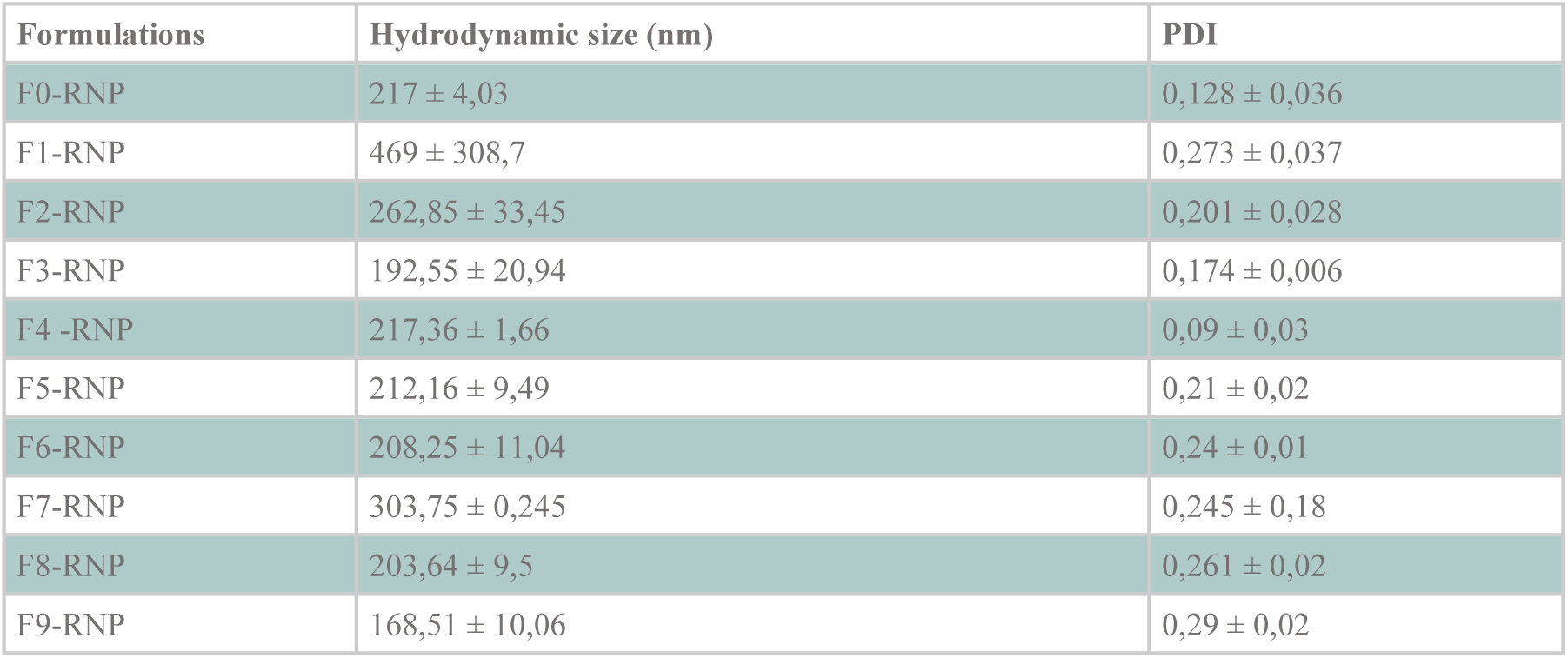
Hydrodynamic size (nm) and polydispersity index (PDI) of the LNP formulations measured by DLS (n=1).

| **Formulations** | **Hydrodynamic size (nm)** | **PDI** |
|---|---|---|
| F0-RNP | 217 ± 4,03 | 0,128 ± 0,036 |
| F1-RNP | 469 ± 308,7 | 0,273 ± 0,037 |
| F2-RNP | 262,85 ± 33,45 | 0,201 ± 0,028 |
| F3-RNP | 192,55 ± 20,94 | 0,174 ± 0,006 |
| F4 -RNP | 217,36 ± 1,66 | 0,09 ± 0,03 |
| F5-RNP | 212,16 ± 9,49 | 0,21 ± 0,02 |
| F6-RNP | 208,25 ± 11,04 | 0,24 ± 0,01 |
| F7-RNP | 303,75 ± 0,245 | 0,245 ± 0,18 |
| F8-RNP | 203,64 ± 9,5 | 0,261 ± 0,02 |
| F9-RNP | 168,51 ± 10,06 | 0,29 ± 0,02 |

It has previously been shown that an excess of oligonucleotides in the formulation increases Cas9 RNP encapsulation into LNPs^20,23^. Therefore, we assessed the effect of increasing the amount of crRNA, by varying the RNP:crRNA ratio, on the colloidal stability and knockout activity of LNP–Cas13b RNP nanocomplexes (Figure S3). Although increasing the crRNA amount slightly reduced LNP size (Figure S3A), it did not improve knockdown efficiency (Figure S3B). Therefore, an RNP:crRNA ratio of 1:2 was selected for subsequent experiments, as higher crRNA amounts did not provide an additional benefit in knockdown efficiency.

Finally, the stability of the LNPs was assessed. LNP–Cas13b RNP samples were stored at 4 °C for 168 h, and colloidal stability (Table 3) and knockdown efficiency (Figure 3) were measured over time. Both colloidal stability and activity remained stable throughout the storage period, demonstrating the high stability of the LNP–Cas13b RNP nanocomplexes.

**Figure 3.**
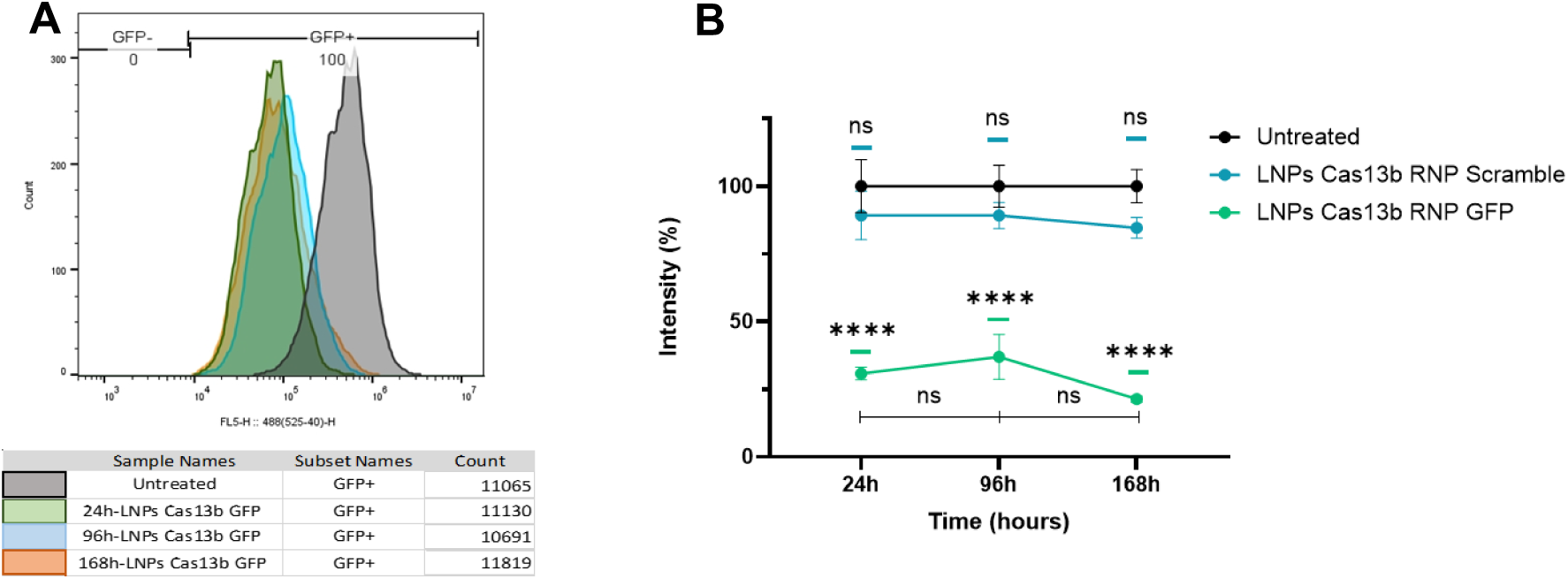
Knock down activity of Cas13b encapsulated into LNPs over time. (A) Representative flow cytometry histograms showing GFP fluorescence intensity (FLS-H:488(525-40)-H) in HeLa cells. The gating strategy differentiates GFP-negative (GFP−) and GFP-positive (GFP+) populations. The table below lists the sample names and cell counts corresponding to each experimental condition: untreated HeLa GFP cells (Untreated), and cells treated with LNPs Cas13b RNPs at different time points (24h, 96h, 168h). (B) HeLa-GFP fluorescence intensity measured by flow cytometry. Comparison of untreated cells, and cells treated with LNPs encapsulating Cas13b RNPs ensembled with a scramble or a GFP specific crRNA. Data are presented as mean ± SD from three independent biological replicates (n = 3). Statistical differences were determined by two-way ANOVA in GraphPad Prism. Brackets indicate the groups compared. ns, not significant (P > 0.05); *, P ≤ 0.05; **, P ≤ 0.01; ***, P ≤ 0.001; ****, P ≤ 0.0001.

**Table 3.** Hydrodynamic size (nm) and polydispersity index (PDI) of the F9 formulation encapsulating Cas13b RNPs over time measured by DLS (n=3).

| **Conditions** | **Time (hours)** | **Hydrodynamic size (nm)** | **PDI** |
|---|---|---|---|
| LNPs Cas13b RNP Scramble | 24 | 156,87 ± 1,71 | 0,140 ± 0,016 |
|  | 96 | 150,60 ± 0,58 | 0,100 ± 0,008 |
|  | 168 | 153,50 ± 3,65 | 0,216 ± 0,022 |
| LNPs Cas13b RNP GFP | 24 | 166,43 ± 3,89 | 0,285 ± 0,024 |
|  | 96 | 148,80 ± 1,48 | 0,110 ± 0,011 |
|  | 168 | 155,17 ± 1,91 | 0,197 ± 0,017 |

### 2.3. Cas13b RNP-mediated knock down of Survivin in pancreatic cancer cell lines

Survivin is involved in cancer cell survival and is overexpressed in chemotherapy-resistant cells. To assess this in pancreatic cancer cell lines, we determined survivin expression in three pancreatic cancer cell lines and compared it with that in non-tumoral cells (Figure S4). Whereas BxPC3 and MIA-PaCa-2 cells showed survivin expression levels similar to those of the non-tumoral cell line, PANC-1 cells clearly overexpressed survivin, as previously described^24^ and in agreement with the gemcitabine-resistant phenotype of this cell line.

Three PspCas13b guide RNAs targeting BIRC5 (survivin) were designed and tested for BIRC5 knockdown in PANC-1 cells by Lipofectamine-mediated transfection (Figure S5). All three guides reduced survivin expression, although with different efficiencies; guide C was therefore selected for subsequent experiments. LNPs encapsulating Cas13b RNPs containing guide C were generated (Table 4), and an encapsulation efficiency of 80% was achieved (Figure 4A).

**Figure 4.**
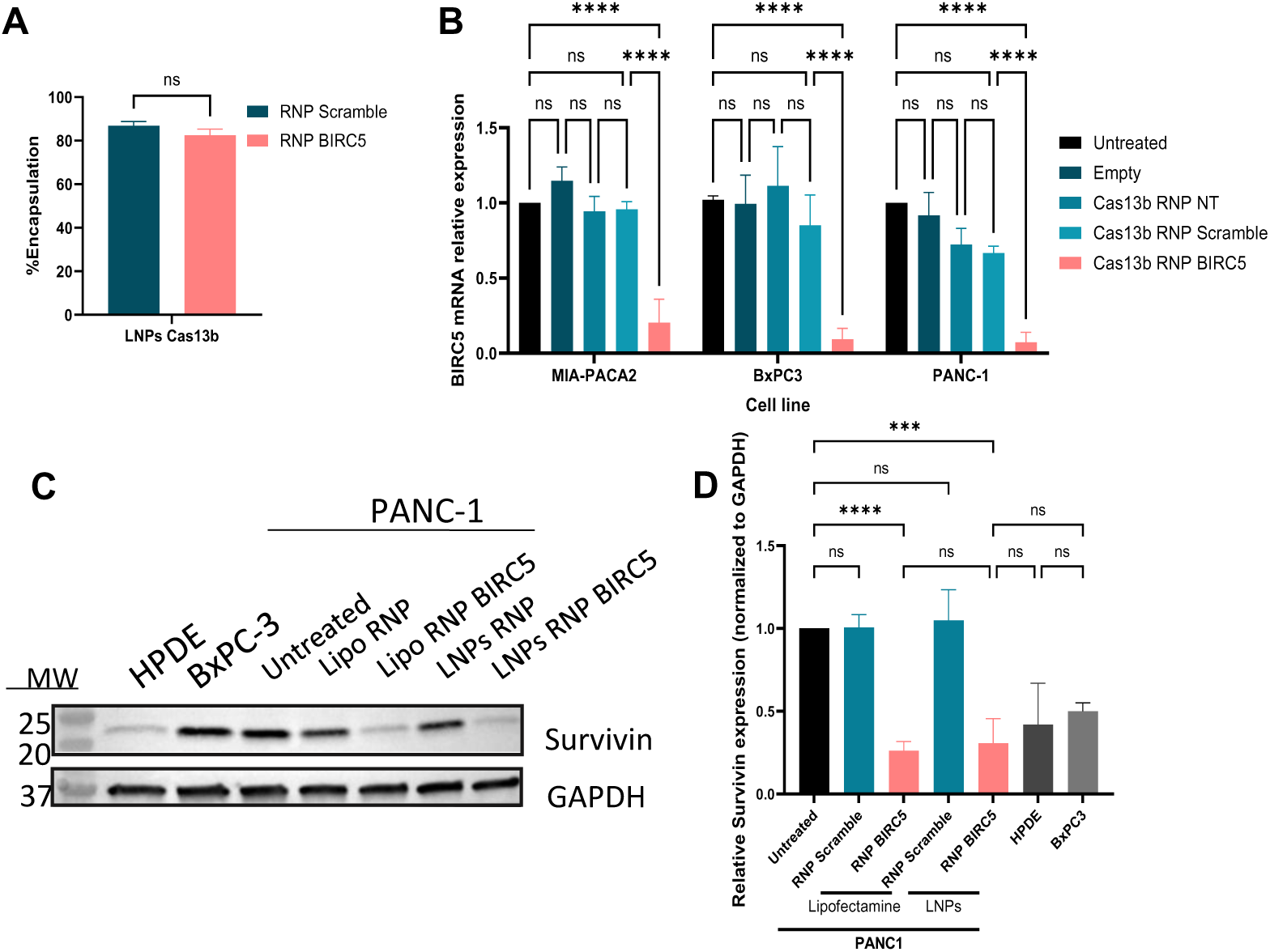
BIRC5 knockdown mediated by LNP-encapsulated Cas13b RNPs. (A) Encapsulation efficiency of Cas13b ribonucleoprotein (RNP) complexes containing either a scrambled guide (RNP Scramble) or a BIRC5-targeting guide (RNP BIRC5) in lipid nanoparticles (LNPs). (B) Relative BIRC5 mRNA expression measured by RT-qPCR in MIA-PaCa2, BxPC3, and PANC-1 cells following treatment with empty LNPs or LNPs encapsulating Cas13b RNPs, as indicated. (C) Representative western blots showing survivin and GAPDH protein levels in untreated PANC-1 cells and PANC-1 cells treated with scrambled or BIRC5-targeting Cas13b RNPs delivered using Lipofectamine or LNPs. HPDE and BxPC3 cells were included for comparison. (D) Densitometric quantification of the western blot signals using ImageJ. Survivin protein levels were normalized to GAPDH and expressed relative to untreated PANC-1 cells. Data are presented as mean ± SD from three independent biological replicates (n = 3). Statistical differences were assessed by one-way ANOVA in GraphPad Prism. Brackets indicate the groups compared. ns, not significant (P > 0.05); ***, P ≤ 0.001; ****, P ≤ 0.0001.

**Table 4.** Hydrodynamic size (nm) and polydispersity index (PDI) of the LNP formulations used for Survivin knockdown measured by DLS (n=7).

| **LNPs Cargo** | **Hydrodynamic size (nm)** | **PDI** |
|---|---|---|
| No Cargo | 155,95 ± 3,13 | 0,13 ± 0,01 |
| Cas13b NT | 166,34 ± 4,21 | 0,21 ± 0,02 |
| Cas13b RNP Scramble | 187,25 ± 8,10 | 0,34 ± 0,01 |
| Cas13b RNP BIRC5 | 161,63 ± 5,13 | 0,26 ± 0,01 |

LNP-delivered Cas13b RNPs reduced survivin expression in all three pancreatic cancer cell lines (Figure 4B). The knockdown efficiency obtained with LNPs was comparable to that achieved with the commercial transfection reagent Lipofectamine (Figure S5B). Western blot analysis in PANC-1 cells, which overexpress survivin, showed that treatment with LNP-encapsulated Cas13b RNPs reduced survivin expression to levels similar to those observed in non-tumoral HPDE cells or in the non-chemoresistant BxPC3 cell line (Figure 4C,D).

Because survivin promotes cell survival, we next assessed whether survivin knockdown induced apoptosis (Figure 5 and Figure S6). Annexin V/DAPI staining followed by flow cytometry showed that LNP-delivered Cas13b RNP treatment increased apoptosis in all three pancreatic cancer cell lines.

**Figure 5.**
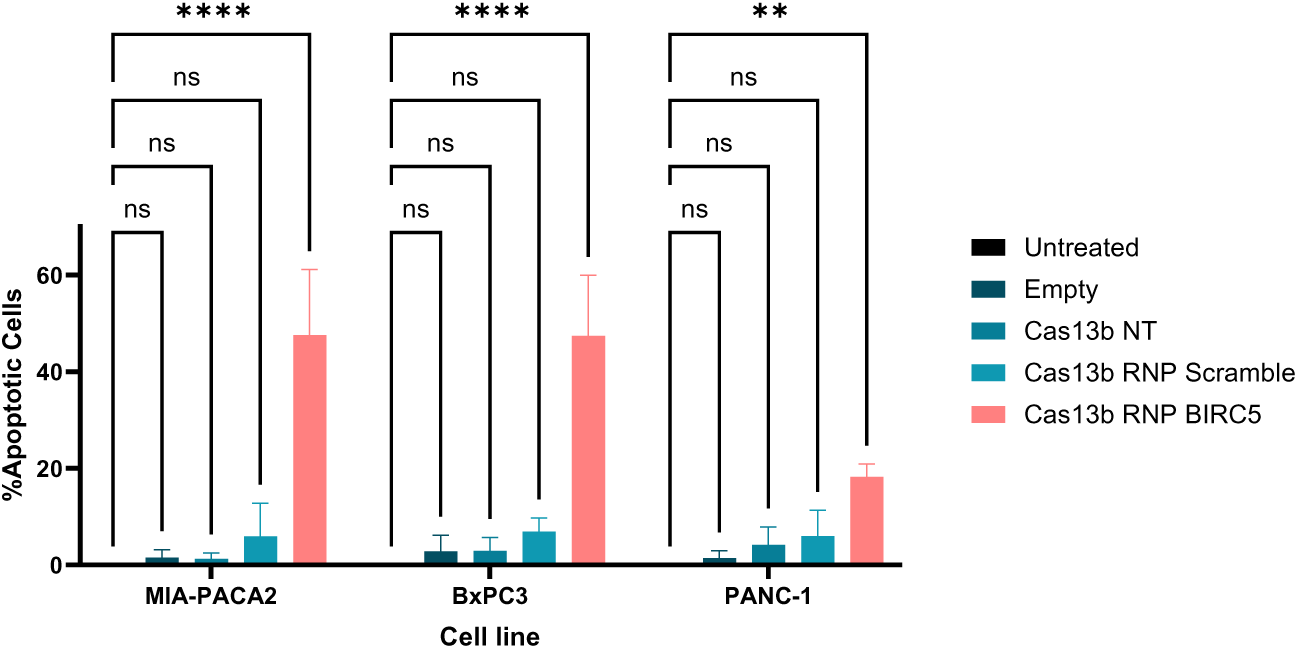
Cas13b RNP-LNPs targeting BIRC5 induce apoptosis in pancreatic cancer cells. Percentage of apoptotic cells (Annexin V⁺) in MIA PaCa-2, BxPC-3 and PANC-1 cells, determined by Annexin V/DAPI flow cytometry, 72 h after treatment. Data are presented as mean ± SD from three independent biological replicates (n = 3). Statistical differences were assessed by one-way ANOVA in GraphPad Prism. Brackets indicate the groups compared. ns, not significant (P > 0.05); **, P ≤ 0.01; ****, P ≤ 0.0001.

Although it has been reported that Cas13b possesses a limited collateral RNase activity in eukaryotic cells^22^, this unspecific and toxic activity is produced upon target recognition and it is therefore highly dependent on target concentration and even cell types^25^. To determine whether the toxicity observed after Cas13b treatment was exclusively due to specific BIRC5 knockdown or also involved collateral activity, total RNA was extracted from treated PANC-1 cells and analyzed using a Bioanalyzer. RNA from treated cells showed integrity comparable to that of untreated samples, indicating the absence of nonspecific RNase activity (Figure S7A). In addition, the integrity of RNA from the housekeeping gene RPL19 was assessed by RT-qPCR after treatment in PANC-1, BxPC3, and MIA-PaCa-2 cells, and no reduction in RNA abundance was observed (Figure S7B). Together, these experiments indicate that, when targeting BIRC5, PspCas13b does not induce nonspecific collateral RNase activity. To further confirm that PspCas13b RNPs lack nonspecific collateral activity even when targeting a highly expressed gene, HeLa cells constitutively expressing GFP were transfected with Cas13b RNPs targeting either GFP or BIRC5. RT-qPCR analysis showed that targeting GFP did not reduce BIRC5 mRNA levels (Figure S7C), and targeting GFP did not reduce GFP fluorescence (Figure S7D). Accordingly, apoptosis was observed only upon specific BIRC5 knockdown and not upon GFP targeting (Figure S7E).

Altogether, these data indicate that the apoptotic response induced by LNP-delivered PspCas13b RNPs is specifically mediated by BIRC5 knockdown rather than by collateral RNase activity.

### 2.4. Synergistic antitumor effect of Survivin knockdown and gemcitabine

The reduction in survivin expression promoted apoptosis and could enhance the cytotoxic effect of gemcitabine, particularly in gemcitabine-resistant tumors. Therefore, we assessed the synergistic effect of the combined treatment in one gemcitabine-sensitive pancreatic cancer cell lines (BxPC3) and one gemcitabine-resistant cell line (PANC-1) (Figure 6). A synergistic effect between both treatments was observed at gemcitabine concentrations of 1–10 µM and Cas13b RNP concentrations of 25–50 nM, but only when the crRNA specifically targeting BIRC5 was used.

**Figure 6.**
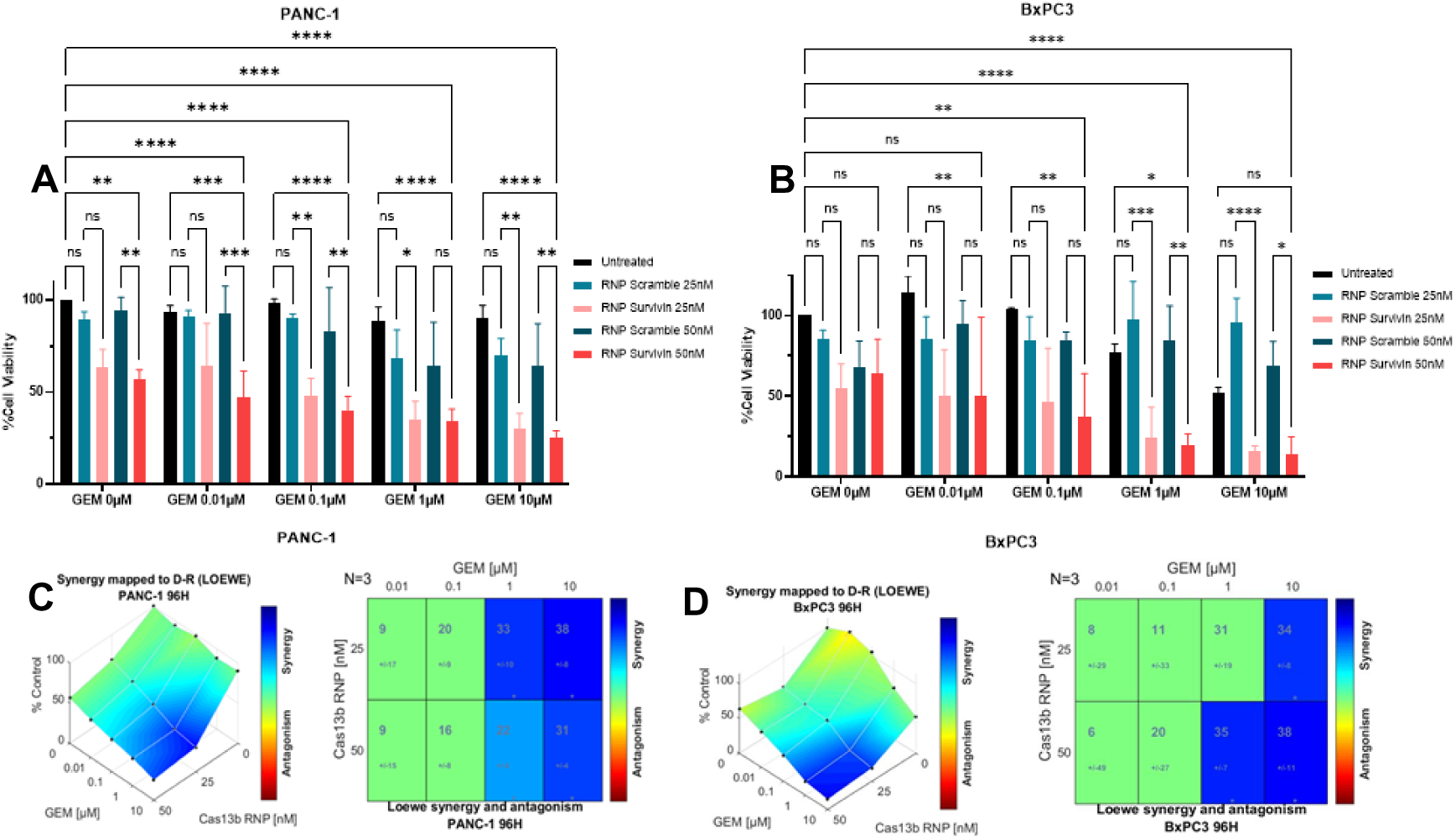
Synergistic effect of LNPs-Cas13b RNP and gemcitabine. (A, B) Cell viability of PANC-1 (A) and BxPC-3 (B) cells after 96 h of treatment first with LNPs loaded with Cas13b RNP and a scrambled crRNA (RNP Scramble) or a BIRC5-targeting crRNA (RNP Survivin) at 25 or 50 nM and 48h later with increasing concentrations of gemcitabine (GEM; 0, 0.01, 0.1, 1 and 10 µM) alone or combined with the LNPs. Cell viability was determined using resazurin (alamarBlue assay) and is expressed as a percentage relative to untreated cells (100% viability), after subtracting the background signal of resazurin in cell-free wells. Data are presented as mean ± SD from three independent biological replicates (n = 3). Statistical differences were assessed by one-way ANOVA in GraphPad Prism. Brackets indicate the groups compared. ns, not significant (P > 0.05); *, P ≤ 0.05; **, P ≤ 0.01; ***, P ≤ 0.001; ****, P ≤ 0.0001. (C, D) Drug interaction analysis for PANC-1 (C) and BxPC-3 (D) cells. Left, three-dimensional dose–response surfaces of cell viability (% of control) mapped to the Loewe additivity model. Right, synergy matrices showing the Loewe synergy/antagonism score for each combination of GEM and Cas13b RNP Survivin (n = 3). Positive values (blue) indicate synergy and negative values (red) indicate antagonism. Synergy analysis and the associated statistics were performed with Combenefit software under the Loewe model (*, P ≤ 0.05).

Next, we evaluated the combinatorial antitumor effect in a xenograft model of gemcitabine-resistant pancreatic cancer. PANC-1 xenografts were established in nude mice and treated intratumorally with LNP-encapsulated Cas13b RNPs targeting BIRC5 and/or gemcitabine. LNPs containing a scrambled Cas13b RNP were included to assess the effect of intratumoral LNP injection alone. Figure 7 shows that only the combination of LNP-delivered Cas13b RNPs targeting BIRC5 and gemcitabine inhibited tumor growth, while the treatment is not toxic (Figure 7D).

**Figure 7.**
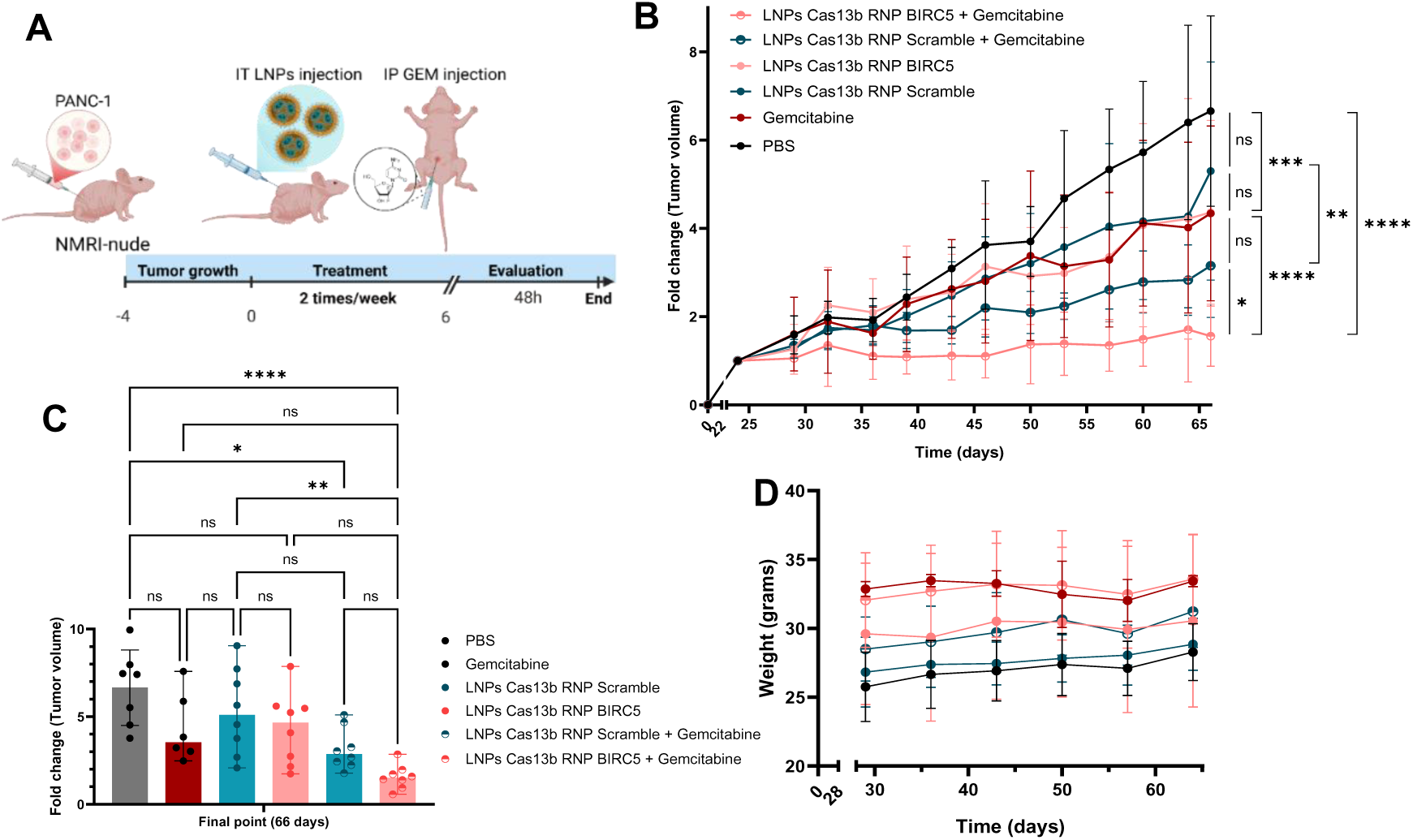
In vivo anti-tumor effect of LNPs-Cas13b RNP and gemcitabine. A) Scheme of the Tumor generation and treatment administration through Intratumoral injection in the case of the LNPs or intraperitoneal injection in the case of the gemcitabine. B)Fold change in tumor volume (mm3) evaluated throught time. C) Fold change in tumor volume at final point (66 days). D) Weight of the mice throught time and the different treatment groups. Data are presented as mean ± SD from at least seven independent biological replicates (n = 7). Statistical differences were assessed by two-way (B) or one-way ANOVA (C) in GraphPad Prism. Brackets indicate the groups compared. ns, not significant (P > 0.05); *, P ≤ 0.05; **, P ≤ 0.01; ***, P ≤ 0.001; ****, P ≤ 0.0001.

To further determine whether this tumor growth inhibition was mediated by the specific activity of Cas13b, tumor xenografts were treated with DiD-labeled LNP–Cas13b RNPs to assess biodistribution and transfection efficiency (Figures 8 and 9A), and BIRC5 knockdown was evaluated by RT-qPCR (Figure 9B). Intratumoral administration confined the LNPs to the tumor and prevented their distribution to other organs, including the liver (Figure 8), thereby minimizing potential off-target effects in non-tumor tissues. Within the tumor, 8.6% of cells were transfected, with high variability (4-15%) (Figure 9A). Furthermore, BIRC5 expression was analyzed in whole-tumor samples after the different treatments (Figure 9B). As previously reported, gemcitabine treatment increased BIRC5 expression^10^, whereas its combination with BIRC5-targeting Cas13b RNPs reduced expression to near-basal levels. A clear reduction in BIRC5 expression was also observed in tumors treated with BIRC5-targeting RNPs compared with those treated with scrambled RNPs; however, neither difference reached statistical significance, likely owing to experimental variability (Figure 9B).

**Figure 8.**
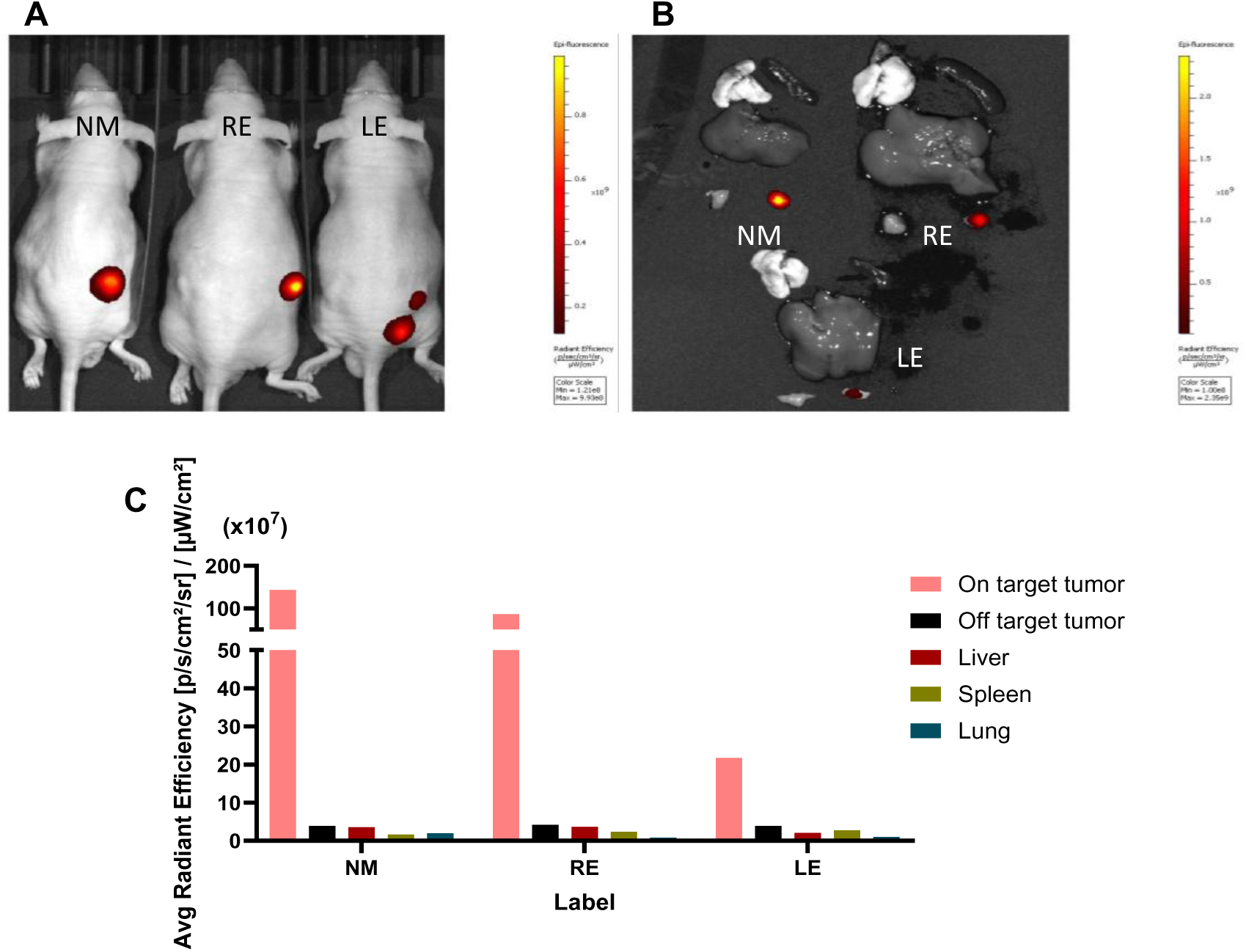
In vivo and ex vivo distribution of DiD-labeled LNPs containing BIRC5-targeting Cas13b RNPs. (A) Representative in vivo fluorescence images of three mice bearing bilateral subcutaneous tumors, with one tumor in each flank. Left-flank tumors were injected with unlabeled LNPs containing BIRC5-targeting Cas13b RNPs, whereas right-flank tumors were injected with the corresponding DiD-labeled formulation. Mice were identified by ear markings: NM, no marking; RE, right ear; and LE, left ear. (B) Ex vivo fluorescence images of tissues collected from the same mice. For each mouse, tissues are arranged as follows: lung (upper left), spleen (upper right), liver (center), left-flank tumor (lower left), and right-flank tumor (lower right). Fluorescence signals in panels A and B are displayed as pseudocolor overlays on the corresponding photographic images. (C) Average radiant efficiency of the DiD signal measured ex vivo in the injected tumor (on-target tumor), contralateral tumor (off-target tumor), liver, spleen, and lung of each mouse

**Figure 9.**
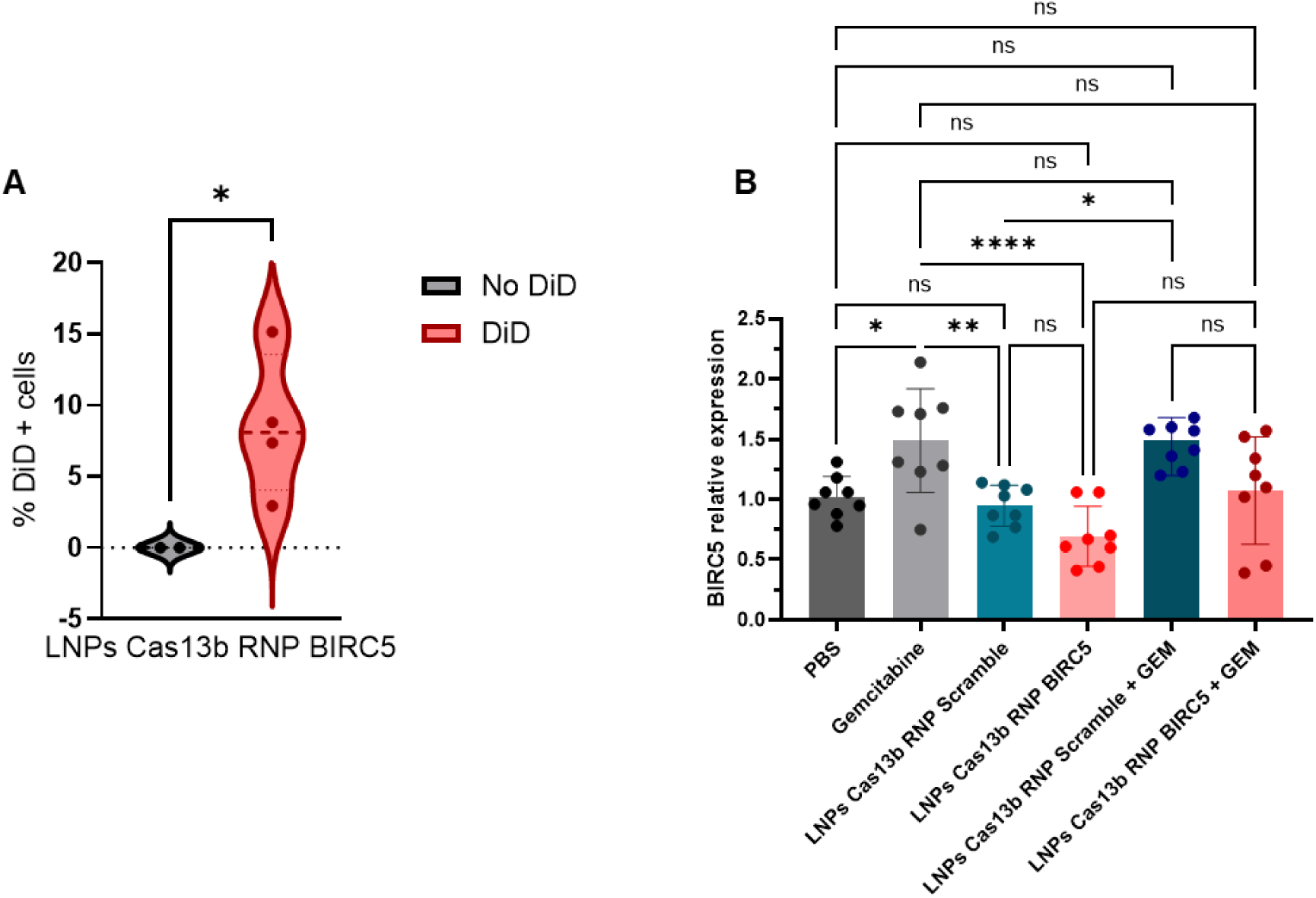
Evaluation of specific activity of Cas13b inside tumors, A) Percentage of transfected cells as percentage of DiD positive cells inside the tumors treated with DiD positive LNPs. Data is presented as mean ± SD from four independent biological replicates (n = 4). Statistical differences were assessed by one-way ANOVA in GraphPad Prism. Brackets indicate the groups compared. ns, not significant (P > 0.05); *, P ≤ 0.05. B) BIRC5 mRNA relative expression measured by RT-qPCR of the tumors extracted from each treatment group 48h after the treatment. Data is presented as mean ± SD from eight independent biological replicates (n = 8). Statistical differences were assessed by one-way ANOVA in GraphPad Prism. Brackets indicate the groups compared. ns, not significant (P > 0.05); *, P ≤ 0.05; **, P≤ 0.01; ****, P ≤ 0.0001.

## 3. Discussion

In this study we show that pspCas13b ribonucleoproteins (RNPs) can be encapsulated in lipid nanoparticles (LNPs) and delivered into cells to silence endogenous transcripts, and that LNP-Cas13b RNPs targeting BIRC5 (survivin) sensitize gemcitabine-resistant pancreatic cancer cells in vitro and in vivo. Most Cas13 applications in mammalian cells rely on plasmids, viral vectors or mRNA that must be transcribed and/or translated inside the cell^12,26^. RNPs bypass these steps. They are active as soon as they reach the cytosol, carry no risk of genomic integration, and are cleared within a limited time. For Cas9, RNP delivery achieved efficient editing with fewer off-target events than plasmid expression^18,27^. This rationale is arguably stronger for Cas13, whose collateral RNase activity and guide-independent toxicity scale with the abundance and persistence of the effector^17,22^. Sustained Cas13 expression has been toxic in primary neurons, zebrafish embryos, flies and adult mice^16,28,29^, and regulatory circuits have been needed to diminish it^30^. A transient, dose-controlled RNP is therefore an attractive format to widen the therapeutic window of Cas13. To our knowledge, LNP-based delivery of Cas13 has so far been limited to mRNA^31^ or plasmid DNA^32^, and this is the first report of Cas13 RNPs formulated in LNPs.

LNPs are well suited to this cargo. They protect the RNP from degradation, can be produced reproducibly, and can be tuned through the ionizable, helper and PEG lipids^33^. Their main limitation is that, after systemic administration, they are opsonized by apolipoprotein E and taken up mainly by hepatocytes^34^. This hampers delivery to extrahepatic tumors and increases hepatic exposure. Extrahepatic targeting has been explored, mainly by modifying the physicochemical properties of LNPs or by adding targeting molecules (peptides, antibodies, aptamers) to their surface. However, only a limited number of tissues have been targeted liver, kidneys, lungs, bone marrow, and tumors^20,35^, supporting the in-site treatment with LNPs.

Particle size and polydispersity determine uptake, biodistribution and potency^36^, and a PDI below 0.3 is generally considered acceptable for lipid carriers^37^. Only four of ten formulations (F3, F6, F8 and F9) were non-toxic, and only F8 and F9 combined low toxicity with knockdown activity, with F9 being the best performing formulation. F3 lacks DOTAP and F6 contains less of it, in line with the need for a positive charge to retain the RNP. DOTAP, however, was associated with toxicity in other formulations^38^, although not in our F8 and F9 formulations. Strikingly, F4 and F8 have the same molar composition and differ only in the ionizable lipid (C12-200 vs DLin-MC3-DMA), yet F4 is toxic while F8 is not. Thus, the structure of the ionizable lipid, and probably its interaction with a large protein–RNA complex rather than a nucleic acid, strongly conditions toxicity. This should be considered when transferring LNP recipes optimised for siRNA or mRNA to protein-based cargos. Our selected formulation also encapsulates Cas13b RNP efficiently, although its encapsulation was determined indirectly quantifying the crRNA. Interestingly, F9-RNP remained colloidally stable and fully active after 168 h at 4 °C. This contrasts with mRNA-LNPs, which are prone to hydrolysis and lipid-mediated degradation of the mRNA and usually require frozen storage^39^. Since the crRNA is shielded within the protein, RNP-LNPs could ease storage and logistics. Longer storage, freeze–thaw cycles and lyophilisation remain to be tested.

Our pspCas13b RNP showed no evidence of collateral activity. RNA integrity was preserved, RPL19 was unchanged, and targeting GFP neither reduced BIRC5 mRNA nor induced apoptosis. This agrees with comparative work showing that Cas13b has less collateral activity than Cas13a or Cas13d^22^. Collateral activity correlates with target abundance^17^, and our GFP-overexpressing cells were therefore a stringent test. This property would extend Cas13 use to abundant or essential targets, where Cas13d-based tools are problematic.

Survivin is a rational target in pancreatic cancer. It is overexpressed in PANC-1 cells, which are gemcitabine-resistant, and our knockdown brought its levels to those of non-tumoral and non-resistant cells. Survivin contributes to gemcitabine resistance in pancreatic cancer cells^9^, and gemcitabine itself up-regulates survivin, whereas its suppression increases sensitivity^10^. Our data are consistent with this. We observed synergy only with the BIRC5-targeting guide, and only the combined treatment inhibited tumor growth in vivo. The up-regulation of BIRC5 by gemcitabine and its return to basal levels with Cas13b also fit this model, although this effect did not reach statistical significance. Survivin-targeting antisense oligonucleotides reached the clinic but failed because of insufficient and inconsistent target engagement, limited tumor delivery and dose-limiting toxicities^5,11^. Combining a different silencing modality with local administration addresses these two issues.

Intratumoral administration was chosen to maximize the local dose and avoid hepatic and systemic exposure, and DiD-labelled LNPs remained confined to the tumour. Pancreatic ductal adenocarcinoma is poorly vascularised and surrounded by a dense stroma that limits the penetration of systemically administered agents^40^, so local injection is a reasonable option. It is also clinically feasible, since endoscopic ultrasound-guided injection is already used in humans. A siRNA-eluting implant targeting KRASG12D was safe and showed potential efficacy together with gemcitabine in locally advanced disease^41^, and endoscopic intratumoral injection of an oncolytic virus with gemcitabine has also been tested^42^. Nevertheless, intratumoral delivery is no guarantee of efficacy, as shown by the negative phase III trial of intratumoral TNFerade^43^. It is also restricted to accessible, locally advanced or borderline-resectable tumors, so it does not address metastatic disease. The most plausible setting for our approach would therefore be locally advanced disease, for example in a neoadjuvant combination with gemcitabine, while systemic versions with extrahepatic-targeted LNPs^35^ would be needed for disseminated disease.

Despite the limitations of our study, like the variable and not significant intratumoral BIRC5 knockdown, , in vivo mechanism should be reinforced with larger cohorts, protein-level readouts and orthotopic models; and the use of nude mice, that do not allow assessment of immunogenicity^44^, our LNP-formulated Cas13b RNPs combine transient activity, no detectable collateral RNase activity and storage stability, and their combination with gemcitabine overcame drug resistance in a pancreatic tumour model, supporting further development of this platform.

## 4. Materials and Methods

### 4.1. Guide RNA synthesis

All guide RNAs (gRNAs) used in this study were synthesized by in vitro transcription. DNA oligonucleotides complementary to the target gRNA and containing the T7 promoter sequence were purchased from Integrated DNA Technologies (IDT). DNA templates were annealed with the corresponding T7 antisense oligonucleotide at a 1:1.5 molar ratio (DNA template:T7 antisense oligonucleotide) using a Mastercycler thermocycler (Eppendorf). Annealing reactions contained 4 µM DNA template.

In vitro transcription was performed using the MEGAscript T7 Transcription Kit (Thermo Fisher Scientific) according to the manufacturer’s instructions. Briefly, each 20 µL reaction contained 2 µL of each ribonucleotide solution, 2 µL reaction buffer, 2 µL enzyme mix, 6 µL of a 1:10 dilution of the annealed template, and nuclease-free water to the final volume. Reactions were incubated at 37 °C for 16 h. Template DNA was subsequently removed by addition of 1 µL TURBO DNase per reaction, followed by incubation at 37 °C for 30 min.

Transcribed RNAs were analyzed by denaturing 10% polyacrylamide gel electrophoresis containing 8 M urea to assess size and integrity. RNA was purified by phenol–chloroform extraction. Briefly, 115 µL nuclease-free water and 15 µL ammonium acetate stop solution were added to each transcription reaction. Samples were purified following RNA Clean and concentrator-5 kit of ZYMO RESEARCH and eluted 20 µL nuclease-free water. Purified gRNAs were stored at −80 °C. All procedures involving phenol and chloroform were performed in a chemical fume hood.

**Table 5:**
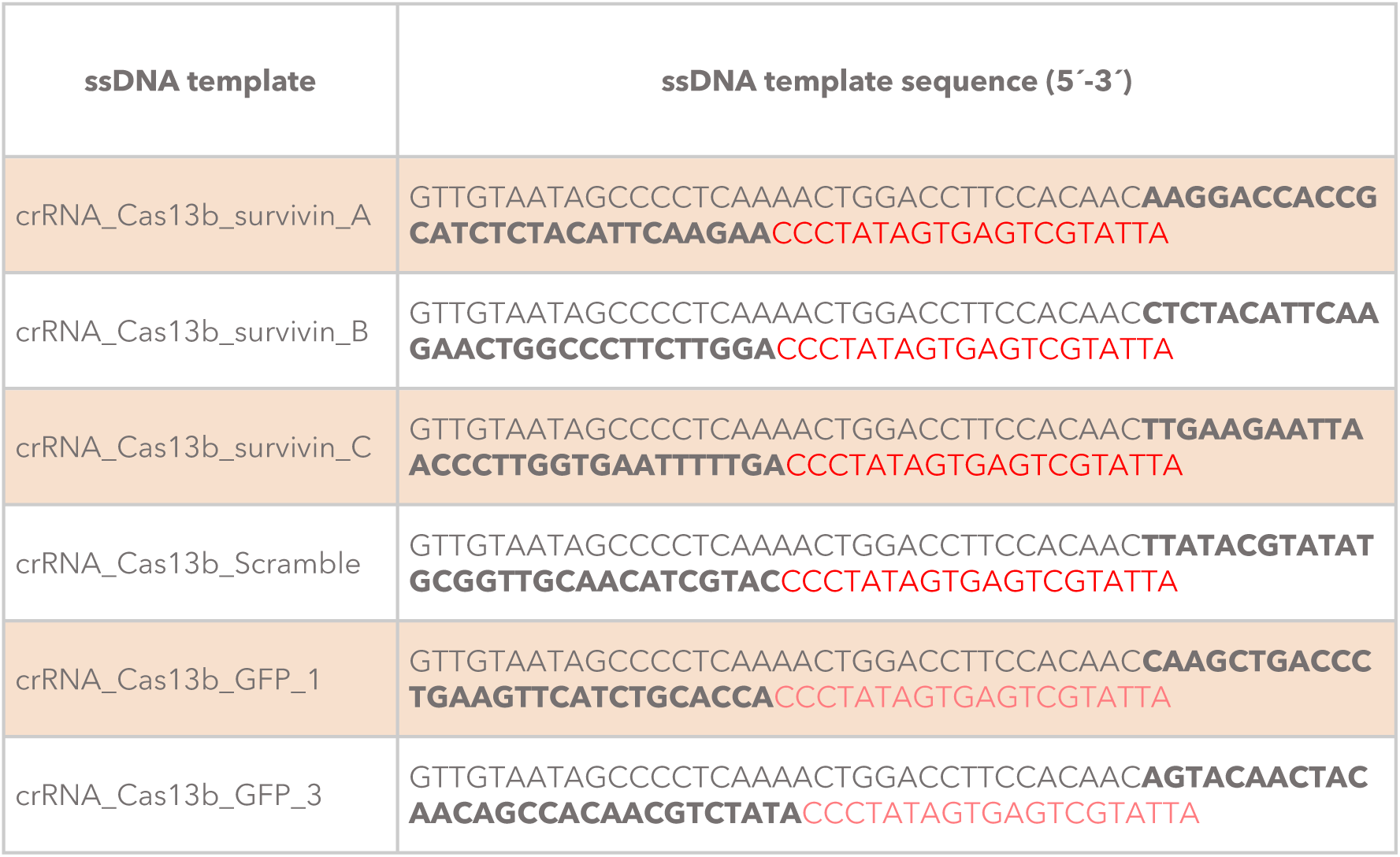
ssDNA templates used for transcription of Cas13 crRNA guides. Regular text represents the direct repeat (DR) region of the crRNA; bold text indicates the spacer region; red text marks the sequence recognized by T7 RNA polymerase.

**Table 6:**
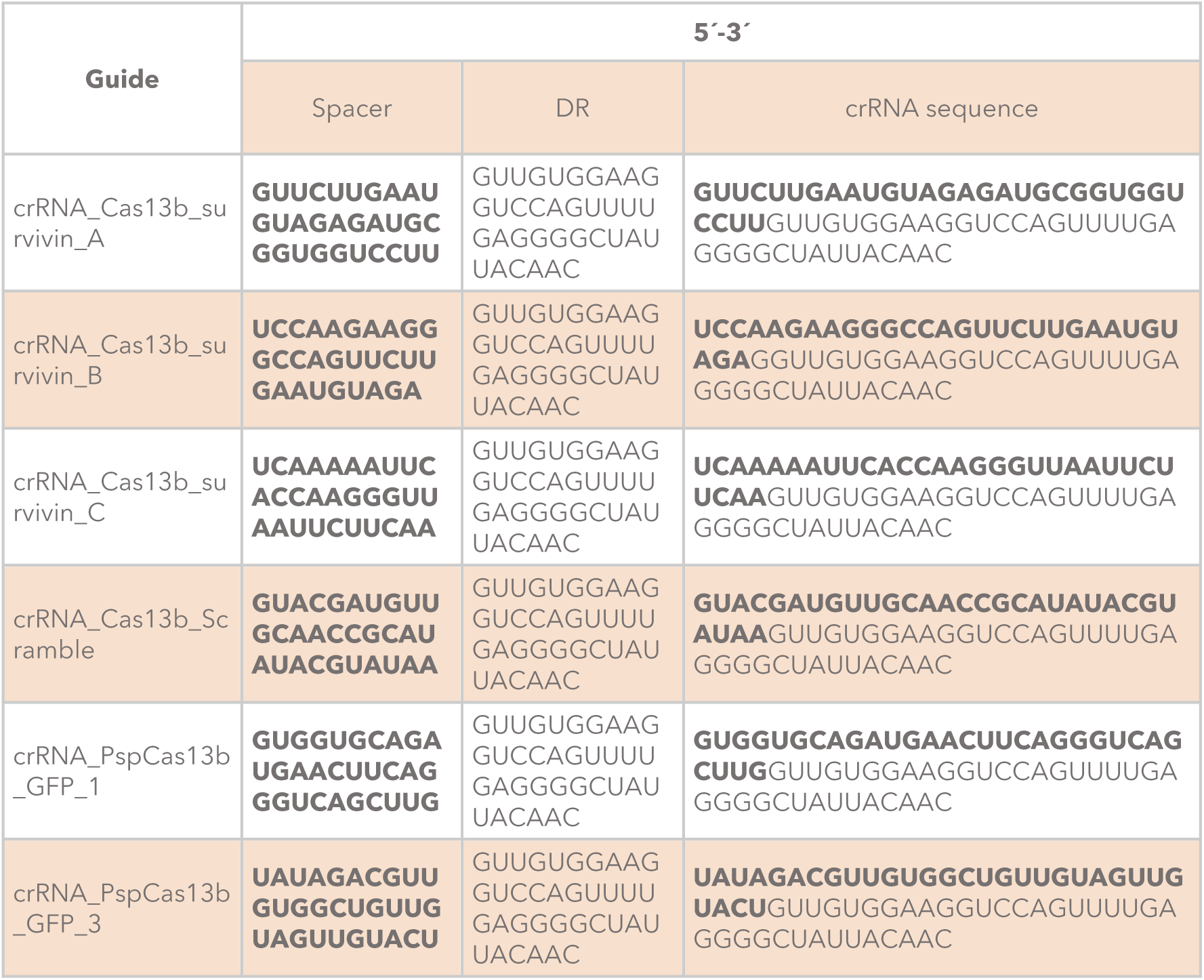
Sequences of crRNA guides for Cas13b.

### 4.3 PspCas13b expression and purification

PspCas13b was expressed from the pC0068-PspCas13b-His6-TwinStrep-SUMO-BsaI plasmid (Addgene plasmid #115219). The plasmid was transformed into chemically competent *Escherichia coli* BL21(DE3) cells. A single colony was used to inoculate 100 mL LB medium supplemented with ampicillin (100 µg/mL), which was incubated overnight at 37 °C and 200 rpm. The starter culture was used to inoculate 3 L LB-ampicillin medium, and bacteria were grown at 37 °C and 200 rpm until OD₆₀₀ reached 0.3. The culture was cooled to 16 °C, and expression was induced at OD₆₀₀ = 0.5–0.6 by addition of 0.5 mM isopropyl β-D-1-thiogalactopyranoside (IPTG). After overnight expression, cells were harvested by centrifugation at 4 °C.

Cell pellets were resuspended in lysis buffer at pH 7.5 containing 50 mM Tris-HCl, 500 mM NaCl, 5% glycerol and supplemented with 5 mM β-mercaptoethanol, 0.5 mM phenylmethylsulfonyl fluoride (PMSF), lysozyme, and EDTA-free protease inhibitor cocktail. Cells were lysed on ice for 45 min followed by sonication using a Model 120 Sonic Dismembrator (Fisher Scientific) with three cycles of 2.5 min at 50% amplitude (10s on/10s off). Lysates were clarified by centrifugation at 11,000 × g for 1 h at 4 °C.

The soluble fraction containing His₆-SUMO-PspCas13b was loaded onto a 5 mL HisTrap HP affinity column (Cytiva) equilibrated with buffer A (50 mM Tris-HCl, 500 mM NaCl, 5% glycerol, 10 mM imidazole, 5 mM β-mercaptoethanol, pH 7.5). After washing with at least five column volumes of buffer A, protein was eluted using a linear gradient from 0% to 100% buffer B (50 mM Tris-HCl, 500 mM NaCl, 5% glycerol, 250 mM imidazole, 5 mM β-mercaptoethanol, pH 7.5). Fractions were analyzed by 10% SDS-PAGE and Coomassie blue staining. Fractions containing PspCas13b were pooled and concentrated using 100-kDa molecular-weight-cutoff centrifugal filters (Merck Millipore).

The His₆-SUMO tag was removed by incubation with ULP1 protease at an enzyme:substrate ratio of 1 mg ULP1 per 5 mg PspCas13b. The reaction was dialyzed overnight at 4 °C against a dialysis buffer at pH 7.5 containing 20 mM Tris-HCl, 500 mM NaCl, 5% glycerol, and 1 mM dithiothreitol (DTT). The sample was clarified by centrifugation at 16,000 × g for 10 min at 4 °C and further purified by size-exclusion chromatography on a Superdex 200 column (Cytiva) equilibrated in the same buffer. PspCas13b-containing fractions were identified by SDS-PAGE, pooled, concentrated using a 100-kDa centrifugal filter, and sterilized through a 0.22 µm syringe filter (Thermo Fisher Scientific).

Protein concentration was determined by measuring absorbance at 280 nm using a NanoDrop One spectrophotometer (Thermo Fisher Scientific) and applying the Beer– Lambert law with the theoretical extinction coefficient (ε=102750). Aliquots were flash-frozen in liquid nitrogen and stored at −80 °C. All purification steps were performed at 4 °C using an ÄKTA Prime Plus FPLC system (Cytiva).

### 4.4 RNP complex formation

PspCas13b was incubated with the corresponding crRNA at Cas13:crRNA molar ratio of 1:2. The 1:1, 1:2, 1:3 molar ratios were tested for LNPs formation experiments. RNP complexes were assembled by mixing Cas13b and crRNA in PBS, using a crRNA volume equal to or lower than the Cas13b volume. Mixtures were incubated at 37 °C for 30 min and kept on ice until use.

### 4.5 LNPs Cas13b RNP production

Lipids were purchased from Avanti Polar Lipids, Cayman Chemical, or MedChemExpress. Lipid stocks were prepared in ethanol, sealed with laboratory film to prevent solvent evaporation, and stored at −20 °C. All lipid manipulations were performed under a chemical fume hood or a laminar-flow cabinet, as appropriate.

#### 4.5.1. LNP formulation

DOPE, cholesterol, DOTAP, and DSPC were prepared at the desired concentrations in ethanol. C12-200, DLin-MC3-DMA, and DMG-PEG were used as supplied in ethanol. Lipid stocks and mixtures were handled using Hamilton glass syringes, which were rinsed sequentially with chloroform and ethanol before and after each use. Lipid mixtures were prepared at 4 mM total lipid concentration unless otherwise stated and stored sealed at −20 °C.

Prior to LNP formation, lipid solutions were warmed to 37 °C in a sonication bath and gently mixed until homogeneous. All formulations intended for cell experiments were prepared under sterile conditions in a laminar-flow cabinet.

#### 4.5.2. LNP-RNP formation

All nanoparticle preparations were performed under sterile conditions in a laminar flow cabinet. For in vitro experiments, PspCas13b ribonucleoprotein (RNP) complexes were preassembled in 1× PBS at a Cas13b:crRNA molar ratio of 1:2. The ethanolic lipid mixture was added to the aqueous RNP solution at a 1:3 (v/v) organic:aqueous phase ratio and gently mixed by pipetting. The total lipid:crRNA mass ratio was approximately 15:1 (w/w). The formulations were incubated for 15 min at room temperature and subsequently kept on ice until use. Empty LNPs were prepared following the same procedure, replacing the RNP solution with 1× PBS. Unless otherwise stated, each well received a 10 µL aliquot containing Cas13b and crRNA at 0.5 and 1 µM, respectively, corresponding to 513.75 ng of Cas13b, 211.7 ng of crRNA and 3180 ng of total lipids (0.5mM) per well for the selected formulation (F9). The molar composition and the amounts of each lipid component are detailed in Table 1.

For in vivo experiments, F9 LNPs were prepared following the same mixing procedure and maintaining a Cas13b:crRNA molar ratio of 1:2. Each tumour received 10 µg of Cas13b, 3.32 µg of crRNA and 26.56 µg of total lipids, corresponding to a lipid:crRNA mass ratio of approximately 8:1 (w/w), following a subsequent optimization of the formulation. After incubation, the formulations were dialysed against 1 L of PBS at 4 °C overnight to remove ethanol and non-encapsulated components.

### 4.6. LNP characterization

#### 4.6.1. Dynamic light scattering

The hydrodynamic diameter and size distribution of LNPs were determined by dynamic light scattering (DLS) using a Litesizer 500 instrument (Anton Paar). LNP samples were diluted 1:10 in 1× PBS. Three independent measurements were acquired per formulation and the number of independent biological replicates is indicated in each case and mean values with SD were reported. DLS is commonly used to determine particle size and distribution, while RNA encapsulation efficiency is commonly quantified using a ribonucleic-acid-binding fluorescent assay.

#### 4.6.2. Encapsulation efficiency

Encapsulation efficiency (EE) was determined using the Qubit RNA High Sensitivity assay kit (Thermo Fisher Scientific). To quantify non-encapsulated crRNA, 1 µL Proteinase K and Qubit reagent were added to intact LNP samples according to the kit protocol. To quantify total crRNA, LNPs were disrupted by addition of 1 µL Tween-20 and 1 µL Proteinase K prior to addition of Qubit reagent. Encapsulation efficiency was calculated as:

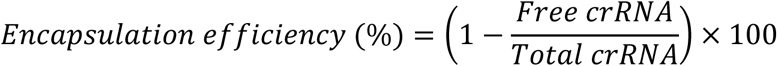

A Cas13b-RNP solution prepared at the expected final crRNA concentration was used as a quantification control.

### 4.7 Mammalian cells transfection

HeLa-WT, HeLa-eGFP, PANC-1, BxPC-3, and MIA-PaCa-2 cells were maintained in DMEM High 10% Fetal Bovine Serum (FBS) and 1% Penicilin/streptamicin at 37 °C in a humidified incubator containing 5% CO₂ (Binder CB210). HPDE was maintained in in Keratinocyte Serum-Free Medium (K-SFM; Gibco, cat. no. 17005042) supplemented with bovine pituitary extract (BPE) and recombinant human epidermal growth factor (EGF) provided by the manufacturer, and with 1% penicillin–streptomycin at 37 °C in a humidified incubator containing 5% CO₂.

Cells were seeded 24 h before transfection to reach approximately 60–70% confluence at the time of treatment. Cells were seeded at 10,000 cells/well in 96-well plates (11,000 cells/well for PANC-1), 50,000 cells/well in 24-well plates (55,000 cells/well for PANC-1), 100,000 cells/well in 12-well plates (110,000 cells/well for PANC-1), and 300,000 cells/well in 6-well plates (330,000 cells/well for PANC-1).

As a positive control were performed using Lipofectamine 2000 (Thermo Fisher Scientific) according to the manufacturer’s instructions with minor modifications. The transfection mixture volume corresponded to 10% of the final well volume. Lipofectamine 2000 was used at 0.3 µL/well in 96-well plates, 1 µL/well in 24-well plates, 1.5 µL/well in 12-well plates, and 2 µL/well in 6-well plates. Cas13 protein or preformed ribonucleoprotein (RNP) complexes were diluted in Opti-MEM I Reduced Serum Medium (Gibco). In parallel, Lipofectamine 2000 was diluted in Opti-MEM. Both mixtures were incubated separately at room temperature for 5 min, combined, and incubated for an additional 20 min. The resulting complexes were added dropwise to the cells.

In the case of LNPs, they were prepared at 100µM lipid concentration, then they were diluted ½ in PBS 1x to reduce exposure to ethanol and added to the plates in different volumes: 10 µL/well in 96-well plates, 50 µL/well in 24-well plates, 100 µL/well in 12-well plates, and 150 µL/well in 6-well plates. Culture medium was replaced 4 h after transfection for both LNPs and Lipofectamine.

### 4.8 Synergy and cytotoxicity

#### 4.8.1. Cell Viability assay

To evaluate both the impact of various LNP lipid formulations on cell viability and the synergistic efficacy of Cas13b RNP-mediated BIRC5 knockdown combined with gemcitabine, cells (Hela-GFP for lipid formulations and BxPC-3 and PANC-1 for the analysis of the synergistic effect) were seeded in 96-well plates and allowed to adhere for 24 h. For the lipid formulation evaluation, cells were treated 24 h post-seeding and assessed after 72 h. Meanwhile, for the synergy assays, cells were first treated with LNP– Cas13b RNP formulations, containing either a BIRC5-targeting or a scrambled control crRNA, at final RNP concentrations of 0, 25, and 50 nM. Forty-eight hours after LNP administration, gemcitabine was introduced across a dose range of 0, 0.01, 0.1, 1, and 10 µM in a matrix combination design.

Cell viability was measured 96 h after gemcitabine administration using a resazurin (Alamar Blue) assay. Briefly, a 1 mg/mL resazurin stock solution in 1× PBS was diluted to a 1% (v/v) working solution in complete culture medium. Cells were incubated with the resazurin working solution for 4 h at 37 °C, and absorbance was measured at 570 nm (reference wavelength 600 nm) using an Infinite 200 Pro microplate reader (Tecan). Cell viability percentage was calculated as:

Viability was calculated as:

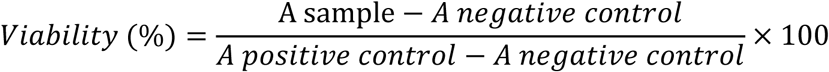

Untreated cells were used as the positive control, whereas complete medium containing 1% (v/v) resazurin but no cells was used as the negative control.

#### 4.8.2. Synergy Quantification and Statistical Analysis

Pharmacological interaction and synergy levels between *BIRC5*-targeting LNP–Cas13b RNPs and gemcitabine were modeled using Combenefit software (version 2.02). Synergy and antagonism scores were calculated based on the Loewe additivity model by mapping dose–response surfaces against expected additive effects. Statistical comparisons between independent treatment groups and controls were performed using GraphPad Prism. Data are expressed as mean ± SD of n = 3 independent biological replicates, and statistical significance was evaluated using two-way ANOVA.

#### 4.8.3. Apoptosis assay

Apoptosis was evaluated 72 h after treatment using Annexin V-FITC and DAPI staining. Cells grown in 24-well or 96-well plates were washed with 1× PBS and resuspended in 100 µL/30 µL 1× Annexin binding buffer (ABB; Thermo Fisher Scientific). For single-stain controls, cells were incubated with either 2 µL /0.4 µL Annexin V-FITC or 1 µL /0.2 µL DAPI (1 µg/µL) in 1× ABB. Samples were incubated for 15 min at room temperature protected from light. Subsequently, 300 µL /50 µL 1× ABB was added to each sample, and fluorescence was acquired using a CytoFLEX flow cytometer (Beckman Coulter). Data was analyzed using FCS Express or FlowJo software.

### 4.9 Assessment of Cas13b activity in mammalian cells

Cis-cleavage activity was assessed in HeLa-WT, HeLa-eGFP, PANC-1, BxPC-3, and MIA-PaCa-2 cells using crRNAs targeting GFP, or BIRC5 mRNAs. Cells were transfected with Cas13b RNPs using Lipofectamine 2000 or the indicated lipid nanocarrier. Knockdown efficiency was evaluated by flow cytometry (72h after treatment), Western blotting (72h after treatment), and/or RT-qPCR (24h after treatment) at the indicated time points.

#### 4.9.1 Flow cytometry

For flow-cytometric analysis, cells cultured in 24-well or 96-well plates were detached, collected by centrifugation at 200 × g for 5 min, washed twice with 500 µL/100 µL PBS, resuspended in PBS, and filtered through 40 µm cell strainers (PluriSelect). Samples were analyzed using CytoFLEX flow cytometer (Beckman Coulter) or Cytomics FC500 flow cytometer (Beckman Coulter) or LSR Fortessa X-20 flow cytometer (BD Biosciences) in the case of Apoptosis samples. Data was analysed using FCS Express or FlowJo software.

#### 4.9.2 Western blotting

Cells were seeded at 300,000 cells/well in 6-well plates and treated 24 h after seeding. At the indicated time points, cells were lysed in 100 µL lysis buffer containing 10 mM Tris-HCl, pH 7.5, 5 mM EDTA, 150 mM NaCl, 10% glycerol, and 0.5% Triton X-100, supplemented with protease and phosphatase inhibitors (50 mM NaF, 1 mM Na₃VO₄, 30 mM sodium pyrophosphate, and 1 mM PMSF). Lysis was performed for 1 h at 4 °C with rotation. Lysates were clarified by centrifugation at 16,000 × g for 15 min at 4 °C, and protein concentration was determined by Bradford assay.

Equal amounts of protein (30–60 µg) were mixed with Laemmli buffer, denatured, and separated by 15% SDS-PAGE under reducing conditions at 150 V. Proteins were transferred onto 0.45 µm nitrocellulose membranes (Cytiva) using a Trans-Blot module (Bio-Rad) in transfer buffer containing 25 mM Tris, 192 mM glycine, and 20% methanol. Membranes were stained with Ponceau S to confirm equal loading and transfer, washed, and blocked with 5% bovine serum albumin (BSA) in TBST (20 mM Tris, 150 mM NaCl, 0.1% Tween-20) for 1 h at room temperature.

Membranes were incubated overnight at 4 °C with primary antibodies diluted in 3% BSA/TBST. The following primary antibodies were used: anti-survivin (D-8; sc-17779; Santa Cruz Biotechnology), anti-GFP (B-2; sc-9996; Santa Cruz Biotechnology), anti-Vinculin (SPM227; ab18058-500; Abcam) and anti-GAPDH (6C5, sc-32233, Santa Cruz Biotechnology). Membranes were washed and incubated for 1 h at room temperature with HRP-conjugated anti-mouse IgG secondary antibody (sc-525409; Santa Cruz Biotechnology). After washing, proteins were detected using Clarity Western ECL Substrate (Bio-Rad), and images were acquired with a ChemiDoc imaging system (Bio-Rad). Band intensities were quantified using Fiji/ImageJ.

#### 4.9.3 RT-qPCR

Cells were seeded at 5 × 10^4^ cells/well in 24-well plates, treated as indicated, and harvested at the specified time points. Total RNA was isolated using the NZY Total RNA Isolation Kit (NZYTech) according to the manufacturer’s instructions and eluted in 30– 40 µL nuclease-free water. RNA concentration was measured using a NanoDrop One spectrophotometer.

Complementary DNA was synthesized from 250–500 ng total RNA using Superscript IV Reverse Transcriptase Kit (Thermo Fisher Scientific) with oligo(dT)₂₃VN primers.

Quantitative PCR was performed using FastStart Universal SYBR Green Master (Rox). Each 20 µL reaction contained 12.5 ng cDNA, 10 µL 2× qPCR master mix/SYBR Green Master, 0.8 µL forward primer (10 µM), 0.8 µL reverse primer (10 µM), and nuclease-free water. Cycling conditions were 95 °C for 10 min, followed by 40 cycles of 95 °C for 15 s and 58 °C for 1 min. Reactions were run in triplicate for each biological replicate using a QuantStudio™ 3 Real-Time PCR System, 96-well, 0.2 mL.

*RPL19*, encoding ribosomal protein L19, was used as the reference gene. Relative gene expression was calculated using the 2^−ΔΔCt^:

Relative expression=2^−ΔΔCt^

where ΔΔCt = ΔCt condition of interest − ΔCt control, and ΔCt = Ct target gene – Ct reference gene.

**Table 7.**
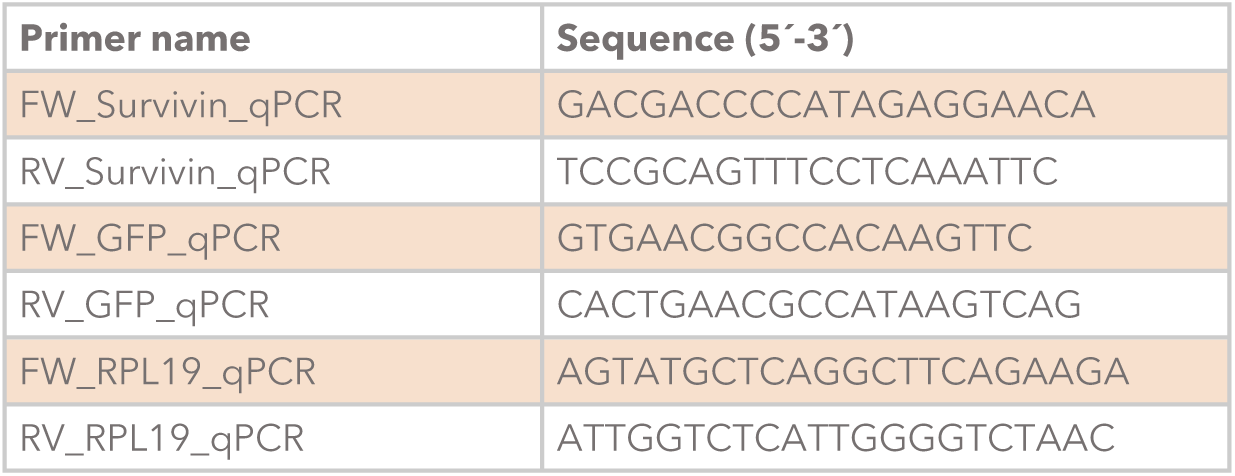
List of primers and their sequences used in this work.

### 4.10. In vivo PANC-1 xenograft model

#### 4.10.1. Cell preparation and xenograft establishment

PANC-1 cells were maintained in high-glucose DMEM supplemented with 10% fetal bovine serum (FBS) at 37 °C in a humidified atmosphere containing 5% CO₂. For xenograft implantation, cells were washed with sterile PBS, detached using trypsin-EDTA, neutralized with complete DMEM, centrifuged at 1,000 × g for 7 min at 20 °C, and resuspended in ice-cold PBS.

Cells were prepared at a density of 5×10^6^ cells per flank. The required cell number was calculated as the number of cells per flank multiplied by the number of mice and by two flanks per mouse. A surplus cell suspension was prepared to account for handling losses. The final suspension was adjusted to deliver 5×10^6^ cells in 50 µL per injection site.

Growth-factor-reduced Matrigel was thawed overnight at 4 °C, and 1 mL syringes were pre-chilled at −20 °C. Immediately before implantation, the cell suspension was mixed 1:1 (v/v) with Matrigel, yielding a final Matrigel concentration of 10 mg/mL. Each injection consisted of 50 µL cell suspension and 50 µL Matrigel, for a total volume of 100 µL per flank. The cell–Matrigel mixture was kept on ice and gently resuspended immediately before loading into chilled syringes.

#### 4.10.2. Animal procedures

All animal procedures were approved by the Animal Welfare and Ethics Committee of CIEMAT and authorized by the competent authority of the Comunidad de Madrid, in accordance with Spanish Royal Decree 53/2013 and EU Directive 2010/63/EU.

Female or male NMRI-nude mice weighing 25–35 g were used. Mice were anesthetized with isoflurane; anesthesia was induced at 3% isoflurane in an induction chamber and maintained at 2.5% isoflurane during implantation, with oxygen supplied at 0.8 L/min.

The cell–Matrigel suspension was loaded into a 1 mL syringe fitted with a 25G needle, taking care to avoid air bubbles. A 100 µL aliquot was injected subcutaneously into each flank. The needle was inserted subcutaneously with the bevel facing upward, and the suspension was administered slowly. The needle was briefly retained in place before withdrawal to minimize reflux. Animals were monitored until complete recovery from anesthesia.

#### 4.10.3. Treatment allocation and dosing

Mice bearing established bilateral tumors were randomly assigned to six experimental groups (n = 4 mice per group). Given that the locally administered LNPs do not diffuse between flanks, each tumor was treated as an independent biological replicate (n = 8 tumors per group). The selected groups were: Untreated, gemcitabine, LNP-Cas13b RNP-Scramble, LNP-Cas13b RNP-BIRC5, LNP-Cas13b RNP-Scramble + Gemcitabine and LNP-Cas13b RNP-BIRC5 + Gemcitabine

The untreated group received sterile physiological saline (0.9% NaCl) as a vehicle control. Mice receiving LNP formulations were administered F9 lipid nanoparticles intratumorally at a total lipid dose of 1.77 mg/kg body weight, corresponding to 53.11 µg lipid per 30 g mouse, or 26.56 µg lipid per tumor. Each tumor received 3.32 µg crRNA and 10 µg Cas13b protein, equivalent to 6.64 µg crRNA (0.221 mg/kg) and 20 µg Cas13b (0.667 mg/kg) per 30 g mouse.

Gemcitabine hydrochloride was dissolved in PBS and administered at 60 mg/kg body weight. Both LNP and gemcitabine treatments were administered twice weekly for 6 weeks.

#### 4.10.4. Tumor monitoring and toxicity assessment

Body weight was recorded once weekly before the second weekly treatment to assess treatment-related toxicity. Tumor formation and growth were monitored twice weekly before treatment administration, beginning after implantation.

Tumor length (L), defined as the longest diameter, and tumor width (W), defined as the shortest perpendicular diameter, were measured using a digital caliper. Tumor volume was calculated using the modified ellipsoid formula:

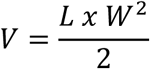

and expressed in mm³. This formula is commonly used for subcutaneous xenograft studies. Tumor volumes are presented as mean ± SD for each experimental group; SEM was calculated as SD/√n. Individual and group-level tumor-growth curves were plotted over time, with sample size indicated for each group. Statistical analyses and graphical representations were generated using GraphPad Prism software.

#### 4.10.5. Tumor total RNA extraction

Tumors were excised, minced, and homogenized in 700 µL QIAzol Lysis Reagent (QIAGEN) using a 5 mm stainless-steel bead and a Mixer Mill MM 400 (RETSCH) at 30 Hz for 4 min. Chloroform (140 µL) was added, samples were vortexed and centrifuged at 13,000 × g for 15 min at 4 °C. The aqueous phase was collected, mixed with 525 µL 100% ethanol, and total RNA was purified using the RNeasy Kit (QIAGEN) according to the manufacturer’s instructions. Relative BIRC5 expression was assessed in extracted total RNA as previously described.

#### 4.10.6. Tumor biodistribution and transfection

Four additional mice bearing tumors (independent of the previous study) were treated with either LNP-Cas13b RNP-BIRC5 or DiD-labeled LNP-Cas13b RNP-BIRC5. F9 lipid nanoparticles were administered intratumorally at a total lipid dose of 1.77 mg/kg body weight, corresponding to 53.11 µg lipid per 30 g mouse and 26.56 µg lipid per tumor. Each tumor received 3.32 µg crRNA and 10 µg Cas13b protein, equivalent to 6.64 µg crRNA (0.221 mg/kg) and 20 µg Cas13b protein (0.667 mg/kg) per 30 g mouse.

For DiD-labeled LNPs, the F9 lipid formulation—composed of DLin-MC3-DMA, DOPE, cholesterol, DMG-PEG, and DOTAP at a molar ratio of 48.9:4.9:37.7:3.7:4.9— was supplemented with DiD at 0.5 mol% relative to total lipid.

Tumor biodistribution of LNPs-Cas13b-DiD was assessed by in vivo and ex vivo fluorescence imaging using an IVIS Spectrum system (PerkinElmer). In vivo images of PANC-1 tumor-bearing mice and ex vivo images of untreated tumors, treated tumors, spleen, liver, and lungs were acquired 16 h after LNPs-Cas13b-DiD administration using Living Image 4.8.0. DiD fluorescence was detected with 660-nm excitation and 710-nm emission filters, a 0.5-s exposure time, an f/2 aperture, and 8 × 8 binning. The field of view was 10 cm for in vivo imaging and 12.5 cm for ex vivo imaging. Reference photographs were acquired with a 0.2-s exposure time, an f/8 aperture, and 2 × 2 binning for anatomical overlay. Images were corrected for background, flat-field, cosmic ray, and CCD artifacts. DiD signal intensity was quantified in counts, and ex vivo organ signals were analyzed using regions of interest (ROIs).

Following imaging, tumors were excised and mechanically minced. Tissue fragments were incubated at 37 °C for 15 min in DMEM containing 100 µg/mL DNase I (Roche, 10104159001), 200 µg/mL Collagenase P (Roche, 11213857001), and 800 µg/mL Dispase II (Sigma-Aldrich, D4693). Samples were vortexed intermittently until no visible tissue fragments remained.

Cell suspensions were filtered through 40 µm Corning cell strainers and centrifuged at 1,200 × g for 5 min. Cell pellets were resuspended in PBS, counted, and adjusted to the appropriate concentration for analysis. DiD-positive cells were quantified using a CytoFLEX flow cytometer (Beckman Coulter), and data were analyzed using FlowJo software.

### Statistical analysis

Statistical analyses were performed using GraphPad Prism (version 10.6.1; GraphPad Software, San Diego, CA, USA) and Combenefit software (version 2.02). Unless otherwise indicated, data are presented as mean ± standard deviation (SD) from at least three independent biological replicates (n = 3). Technical replicates were averaged before statistical analysis when applicable.

The distribution of continuous variables was assessed using the Shapiro–Wilk test, and homogeneity of variances was assessed using the Brown–Forsythe test. Differences among three or more independent groups were analyzed by one-way ANOVA, followed by Šídák’s multiple-comparisons test. Experiments involving two independent factors, such as crRNA specificity and storage time, were analyzed by two-way ANOVA followed by Šídák’s multiple-comparisons test. Comparisons between two groups were performed using an unpaired, two-tailed Student’s t-test.

This approach was applied to LNP hydrodynamic diameter, polydispersity index (PDI), crRNA encapsulation efficiency, GFP and BIRC5 knockdown, apoptosis, and cell viability. Statistical significance was set at P < 0.05. Exact P values are reported in the figures, except where P < 0.0001. In the figures, ns denotes not significant; *, P < 0.05; **, P < 0.01; ***, P < 0.001; and ****, P < 0.0001.

The pharmacological interaction between BIRC5-targeting LNP-Cas13b RNPs and gemcitabine was evaluated using the Loewe additivity model implemented in Combenefit software. Cell viability was measured across a combination matrix of LNP-Cas13b RNP concentrations (0, 25, and 50 nM) and gemcitabine concentrations (0, 0.01, 0.1, 1, and 10 μM). Dose–response surfaces were fitted for each single agent and for the combination, and expected additive effects under the Loewe model were calculated. Synergy scores were derived for each drug combination, with positive values indicating synergy and negative values indicating antagonism. A combination was considered synergistic when the synergy score was positive and statistically significant (P < 0.05), as determined by the statistical module of Combenefit.

In the PANC-1 xenograft study, tumor volume was calculated using the modified ellipsoid formula: V = (L × W²)/2, where L is the longest tumor diameter and W is the shortest perpendicular diameter. Tumor-growth curves were analyzed by two-way repeated-measures ANOVA, followed by Šídák’s multiple-comparisons test. Tumor fold change at the final time point (day 66) was compared among treatment groups using one-way ANOVA followed by Šídák’s multiple-comparisons test. Body weight was analyzed longitudinally using two-way repeated-measures ANOVA. Data are presented as mean ± SD from the number of animals indicated in each figure.

Animals were randomly assigned to treatment groups before treatment initiation. No animals were excluded after randomization. Because intratumoral LNP administration required visual identification of the injection site, investigators were not blinded to treatment allocation. Tumor-volume measurements were performed using a standardized caliper-based protocol.

For biodistribution and intratumoral transfection analyses, DiD fluorescence was quantified as average radiant efficiency in regions of interest corresponding to the injected tumor, contralateral tumor, liver, spleen, and lung. The percentage of DiD-positive cells was determined by flow cytometry. BIRC5 mRNA expression in excised tumors was quantified by RT-qPCR, normalized to RPL19, and expressed using the 2^-ΔΔCt method. These endpoints were compared among treatment groups using one-way ANOVA followed by Šídák’s multiple-comparisons test.

## 6. Author Contributions

DCE: formal analysis, investigation, methodology, visualization, writing-original draft, writing-review and editing. MLV: conceptualization, investigation, methodology, visualization. MS: investigation, resources, methodology. ER: investigation, methodology. LMV: investigation, methodology DP: investigation. PL: resources methodology. AS: supervision, funding acquisition. BS: supervision, funding acquisition, conceptualization, methodology, writing-original draft, writing-review and editing.

