## Supplementary Information for "Lipid nanoparticle-delivered Cas13b ribonucleoproteins targeting survivin overcome gemcitabine resistance in pancreatic cancer"

1. Biomedical Innovation Unit, Center for Research on Energy, Environment and Technology (CIEMAT), Madrid, Spain.
2. Fundación IMDEA Nanociencia, Madrid, Spain.
3. Cellular and Molecular Genitourinary Oncology Group, Institute of Biomedical Research Hospital "12 de Octubre", Madrid, Spain.
4. Tumor Progression Mechanisms Program, Centro de Investigación Biomédica en Red de Cáncer, Madrid, Spain.
5. Department of Chemistry, Biochemistry and Pharmaceutical Sciences,University of Bern, Switzerland
6. Unidad Asociada de Nanobiotecnología (CNB-CSIC e IMDEA Nanociencia) 28049 Madrid Spain.
7. Advanced Therapies Unit, Instituto de Investigación Sanitaria Fundación Jiménez Díaz (IIS-FJD, UAM), Madrid, Spain.
8. Centro de Investigación Biomédica en Red en Enfermedades Raras, Madrid, Spain.

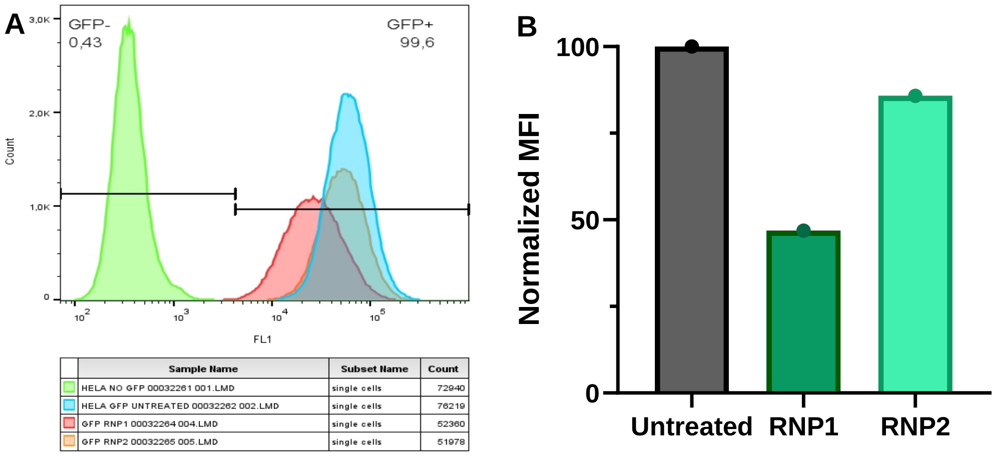


**Figure S1. Flow cytometry evaluation of GFP silencing efficacy mediated by Cas13b RNPs targeting *GFP*. (A)** Representative flow cytometry histograms showing GFP fluorescence intensity (FL1 channel) in HeLa cells. The gating strategy differentiates GFP-negative (GFP−) and GFP-positive (GFP+) populations. The table below lists the sample descriptions and cell counts corresponding to each experimental condition: HeLa cells without GFP (HELA NO GFP), untreated HeLa GFP cells (HELA GFP UNTREATED), and cells treated with two different GFP-targeting Cas13b RNPs (GFP RNP 1 and GFP RNP 2). **(B)** Normalized Mean Fluorescence Intensity (MFI) expressed as a percentage relative to the untreated control group. Data reflect the evaluation of GFP knockdown efficiency for the tested RNP candidates (n=1).

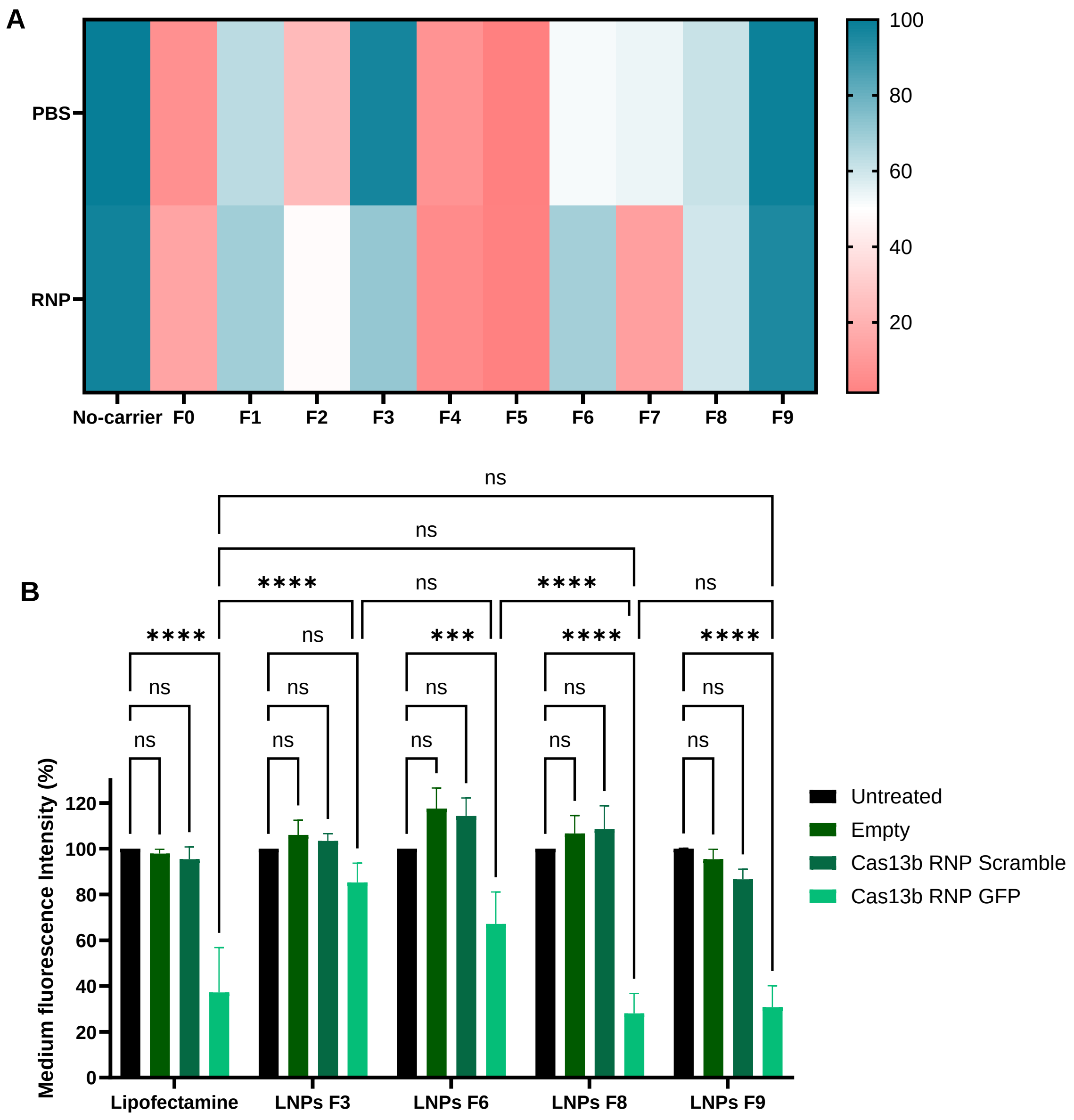


**Figure S2. Screening of lipid formulations for Cas13b RNP delivery.** A) Heatmap showing HeLa-GFP cell viability 72 h after treatment with lipid formulations containing PBS or Cas13b ribonucleoprotein (RNP) complexes, as assessed using a resazurin-based assay (Alamar Blue). Cell viability is expressed as a percentage relative to untreated cells. Data represent the mean of six technical replicates from one biological replicate (n = 1). B) GFP fluorescence in HeLa-GFP cells following treatment with the indicated lipid formulations, assessed by flow cytometry. Median fluorescence intensity is expressed as a percentage relative to untreated cells to evaluate GFP knockdown. Data are presented as the

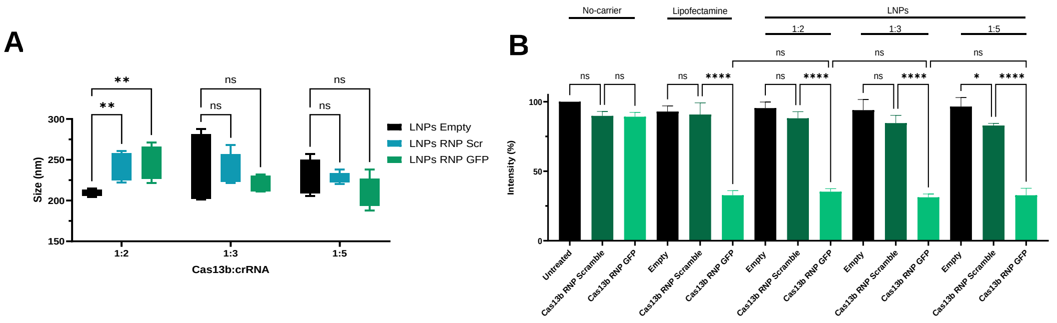


**Figure S3. Optimization of the Cas13b:crRNA ratio for LNP-mediated delivery.** (A) Hydrodynamic diameter of empty LNPs and LNPs encapsulating Cas13b RNPs containing either a scrambled crRNA or a GFP-targeting crRNA, prepared at Cas13b:crRNA ratios of 1:2, 1:3, and 1:5, as measured by dynamic light scattering (DLS). (B) GFP fluorescence in HeLa-GFP cells following treatment with Cas13b RNPs delivered without a carrier, with Lipofectamine, or encapsulated in LNPs prepared at the indicated Cas13b:crRNA ratios, as assessed by flow cytometry. Median fluorescence intensity is expressed as a percentage relative to untreated cells to evaluate GFP knockdown. Data are presented as the mean ± SD of three independent biological replicates (n = 3). Statistical significance was assessed using one-way ANOVA in GraphPad Prism. Brackets indicate the groups compared. ns, not significant (P > 0.05); *, P ≤ 0.05; **, P ≤ 0.01; ****, P ≤ 0.0001.

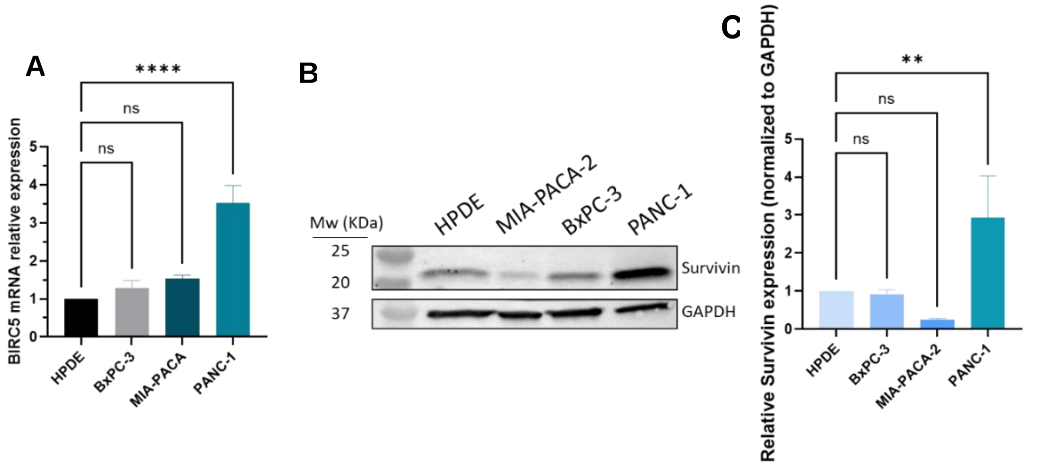


**Figure S4. Basal survivin expression in pancreatic cell lines.** (A) Relative BIRC5 mRNA expression in HPDE, BxPC-3, MIA-PaCa-2, and PANC-1 cells, measured by reverse transcription quantitative PCR (RT-qPCR). (B) Representative western blots showing survivin and GAPDH protein levels in the indicated cell lines. (C) Densitometric quantification of survivin protein levels using ImageJ. Survivin signals were normalized to GAPDH and expressed relative to HPDE cells. Quantitative data are presented as the mean ± SD of three independent biological replicates (n = 3). Statistical significance was assessed

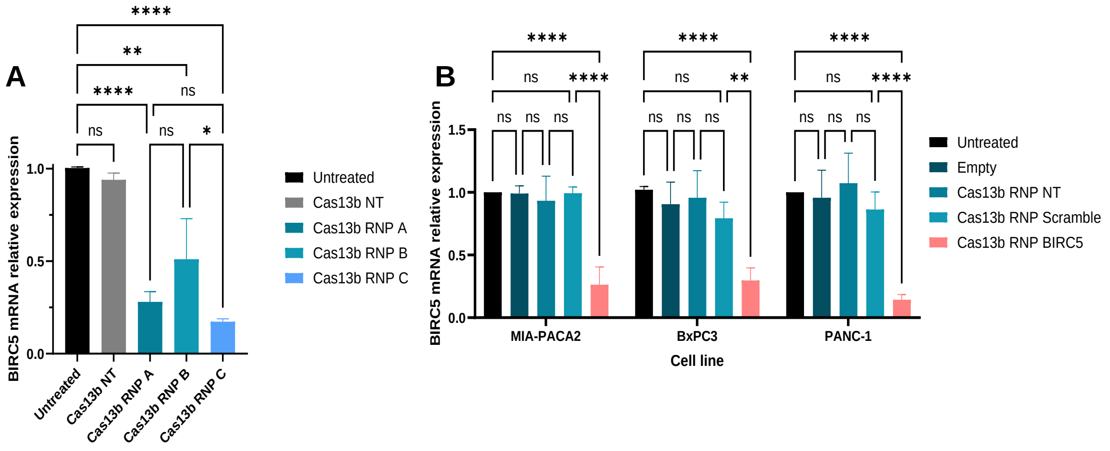


**Figure S5. BIRC5 knockdown mediated by Lipofectamine 2000 transfection of Cas13b RNPs.** (A) Relative BIRC5 mRNA expression measured by RT-qPCR in PANC-1 cells following transfection with non-targeting Cas13b (Cas13b NT) or Cas13b ribonucleoproteins (RNPs) complexed with three different BIRC5-targeting crRNAs (A, B, and C), compared to untreated cells. (B) Relative BIRC5 mRNA expression measured by RT-qPCR in MIA PaCa-2, BxPC-3, and PANC-1 cells treated with Lipofectamine 2000 alone (Empty), Cas13b NT, Cas13b RNP with a scramble guide, or BIRC5-targeting Cas13b RNP, compared to untreated controls. Data are presented as mean ± SD from three independent biological replicates (n = 3). Statistical significance was assessed using one-way ANOVA (A) or two-way ANOVA (B) in GraphPad Prism. Brackets indicate the compared groups. ns, not significant ($P > 0.05$); *, $P \leq 0.05$; , $P \leq 0.01$; ****, $P \leq 0.0001$.

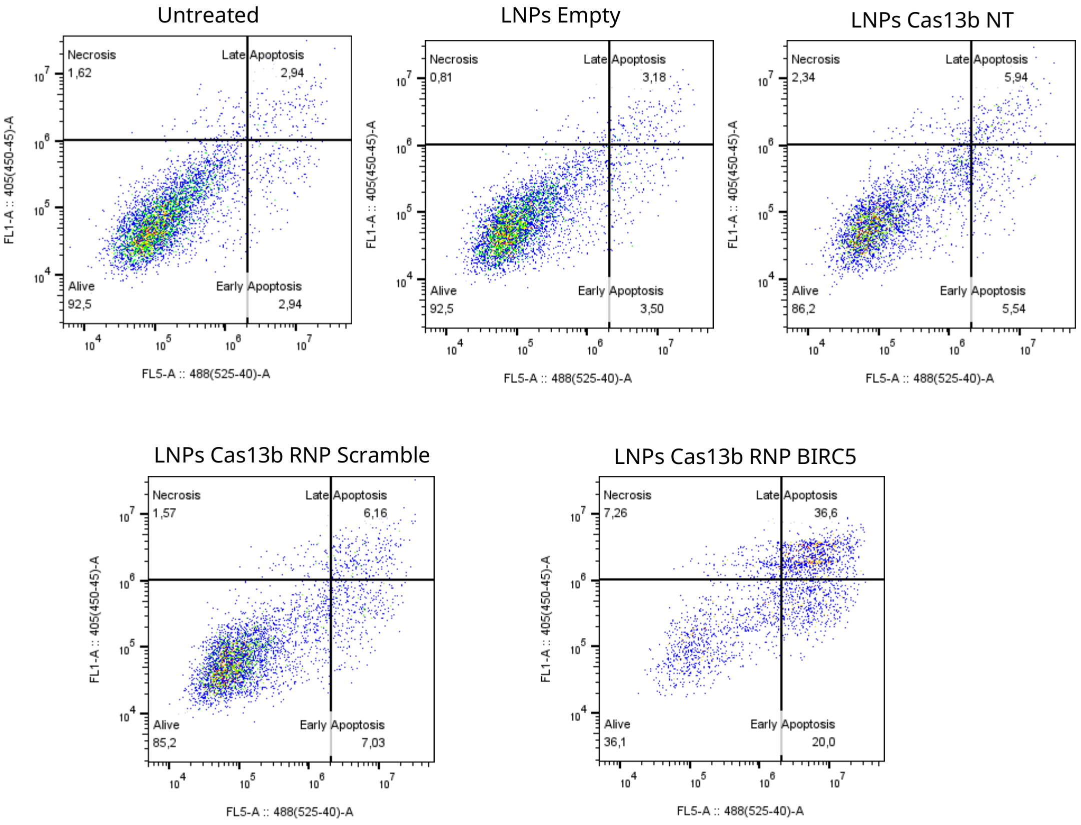

**Figure S6. Representative Annexin V/DAPI flow cytometry analysis of BXPC-3 cells.** Cells were either untreated or treated with empty LNPs, LNPs encapsulating non-targeting Cas13b (Cas13b NT), Cas13b ribonucleoproteins (RNPs) with a scramble guide, or BIRC5-targeting Cas13b RNP. Flow cytometry was performed 72 h post-treatment. Numbers in each quadrant indicate the percentage of viable (Alive), early apoptotic, late apoptotic, and necrotic cells.

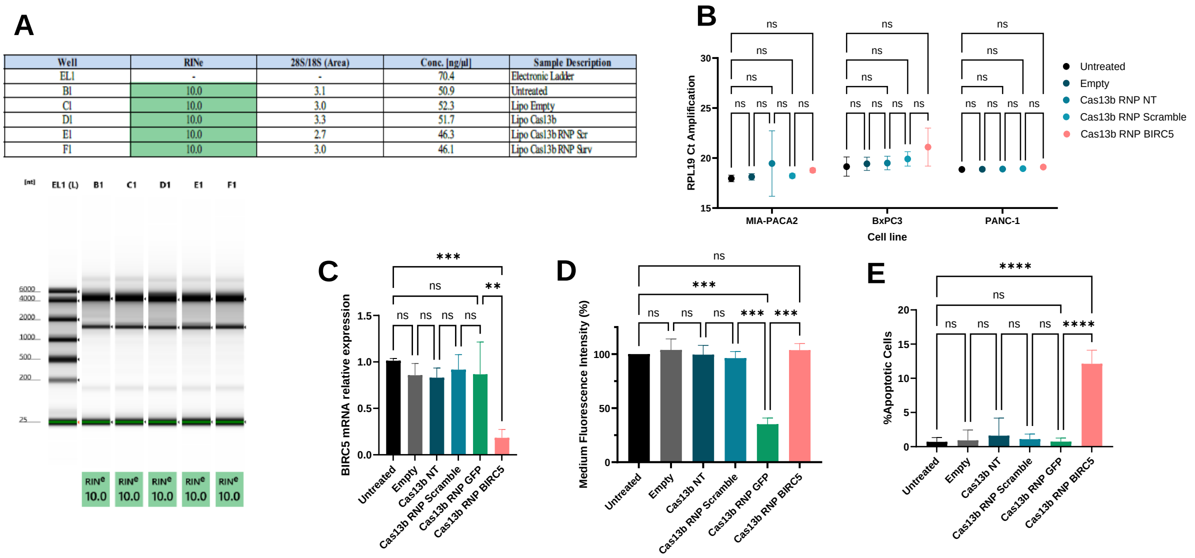


**Figure S7. Evaluation of nonspecific collateral RNase activity produced by pspCas13b RNP transfection.** (A) Total RNA extracted from treated PANC-1 cells and analyzed using a Bioanalyzer. The table (top) and virtual gel (bottom) show the sample description, RNA integrity number equivalent (RINe), 28S/18S area ratio, and concentration for each analyzed well. (B) RT-qPCR amplification Ct values of the housekeeping gene RPL19 in MIA PaCa-2, BxPC-3, and PANC-1 cells following treatment with Cas13b RNPs. (C–E) Assessment of Cas13b RNP trans-activity in HeLa-GFP cells treated with empty LNPs, non-targeting Cas13b, Cas13b RNP with a scramble guide, or Cas13b RNPs targeting GFP or BIRC5, measured by (C) relative BIRC5 mRNA expression via RT-qPCR, (D) mean fluorescence intensity (MFI) expressed as a percentage of the untreated group, and (E) percentage of apoptotic cells evaluated by Annexin V/DAPI flow cytometry. Data are presented as mean ± SD from three independent biological replicates (n = 3). Statistical significance was assessed using one-way (C-E) or two-way ANOVA (B) in GraphPad Prism. Brackets indicate compared groups. ns, not significant ($P > 0.05$); **, $P \leq 0.01$; ***, $P \leq 0.001$; ****, $P \leq 0.0001$.
